# Zebrafish embryonic tissue differentiation is marked by concurrent cell cycle dynamic and gene promoter regulatory changes

**DOI:** 10.1101/2019.12.27.884890

**Authors:** Joseph W Wragg, Leonie Roos, Dunja Vucenovic, Nevena Cvetesic, Boris Lenhard, Ferenc Müller

**Affiliations:** Institute of Cancer and Genomic Sciences, College of Medical and Dental Sciences, University of Birmingham, B15 2TT, Birmingham, UK; Institute of Clinical Sciences and MRC Clinical Sciences Centre, Faculty of Medicine, Imperial College London, London, UK

## Abstract

The core promoter, a stretch of DNA surrounding the transcription start site (TSS) is a major integration point for regulatory signals controlling gene transcription. The process of cell differentiation is accompanied by a marked divergence in transcriptional repertoire between cells of different fates, accompanied by changes in cellular behaviour, in particular their proliferative activity. Investigation of divergent core promoter architectures suggest distinct regulatory networks act on the core promoter, modulating cell behavior through transcriptional profile changes, which ultimately drives key transitions in cellular behaviour during embryonic development. The role that promoter-associated gene regulatory networks play in development associated transitions in cell cycle dynamics (e.g. during differentiation) however, is poorly understood. In this study we demonstrate in a developing *in vivo* model, how core promoter variations play a key role in defining transcriptional output in cells transitioning from a proliferative to cell-lineage specifying phenotype. The FUCCI transgenic system, differentially marks cells in G1 and S/G2/M phases of the cell cycle and can therefore be used to separate rapidly and slowly cycling cells *in vivo*, by virtue of the cell cycle stage they primarily inhabit. Longitudinal assessment of cell proliferation rate during zebrafish embryo development, using this system, revealed a spatial and lineage-specific separation in cell cycling behaviour across post-gastrulation embryos. In order to investigate the role differential promoter usage plays in this process, cap analysis of gene expression (CAGE) was performed on fluorescent associated cell sorted (FACS) FUCCI zebrafish embryos going through somitogenesis, separating cells in accordance with the rate of their cell cycling. This analysis revealed a dramatic increase in lineage and tissue-specific gene expression, concurrent with a slowing of their cell cycling. Core promoters associated with rapidly cycling cells, showed broad distribution of transcription start site usage, featuring positionally constrained CCAAT-box, while slowly cycling cells favoured sharp TSS usage coupled with canonical TATA-box utilisation and enrichment of Sp1 binding sites. These results demonstrate the regulatory role of core promoters in cell cycle-dependent transcription regulation, during somitogenesis stages of embryo development.

## Introduction

The core promoter, a stretch of DNA surrounding the transcription start site (TSS) of all genes, is a key site of transcriptional regulation (1, 2). The advent of 5’ end transcript sequencing (eg. Cap analysis of gene expression [CAGE]), has greatly enhanced our ability to interrogate the role the core promoter of a gene plays in transducing regulatory signals into gene transcription (3-6). Single base pair resolution analysis of TSS location (using CAGE) has revealed an immense diversity in the pattern of transcription initiation on the core promoter, from a narrow distribution of TSSs, with a single base pair dominant site (termed sharp promoters), to a dispersed pattern of TSS usage across the promoter (broad), with a spectrum of different promoter architectures between these two extremes (3-6). Investigation of divergent core promoter architectures have revealed these to be a strong indicator of distinct regulatory networks, acting on the core promoter, modulating cell behavior through transcriptional profile changes (6-9). This is of importance in understanding how key transitions in cellular behaviour during embryonic development are regulated at the level of transcription initiation. The role that promoter-associated gene regulatory networks play in development associated transitions in cell cycle dynamics (e.g. during differentiation) however, is poorly understood.

Embryonic development is marked by several dramatic transitions in the regulatory make up of cells, to permit changes and limitations in their potency, leading to the formation of an organised hierarchical body map. These transitions are often associated with changes in cell cycle dynamics, alongside shifts in transcriptional repertoire (10-13). This process commences with the fusion of two gamete cells into a single fertilised embryo. In many eukaryotes, including zebrafish, this is followed by a number of rapid, synchronous cell cycles, with embryonic behaviour exclusively controlled by maternally deposited factors. At the midblastula transition (MBT) the zygotic genome activates and this process is marked by a slowing of the cell cycle and a loss of synchrony (reviewed in (14, 15). We have previously shown that the transition in cell behaviour from the rapidly cycling synchronous divisions before MBT, to slower, asynchronous, divisions after MBT, accompanied by the activation of the zygotic genome, is marked by a switch in transcription initiation grammar from defined, W-box mediated transcriptional output, to a broader unrestricted initiation grammar, but confined by nucleosome positioning (16). Extensive regulatory reprograming is seen in other model organisms, during this period too, with the first stages of mouse embryo development marked by extensive chromatin remodelling, with lineage-specific expression of several chromatin modifiers, underscoring the potential role of gene regulatory networks in controlling cell fate decisions (17).

Studies using the Fucci cell cycle dynamics reporter in developing zebrafish have revealed that in subsequent stages of development, the process of cell differentiation is marked by a further slowing in cell cycle dynamics as tissue-lineages are specified (18). This is in agreement with *in vitro* studies of human and murine embryonic stem cells, showing that a key indicator of cell differentiation from pluripotency, are transitions in cell cycle dynamics from rapid cycling to a slower cycling identity, characterised by an elongated G1 phase (10-13). In addition to this, studies, investigating transitions from lineage defined multipotent stem cells to terminally differentiated cells, in both muscle (19) and liver development (1), have shown that this process is marked by wholesale depletion of RNA polymerase II regulatory cofactors. Additionally, various cell lineages on the path towards differentiation, appear to be regulated by distinct general transcription factors, forming preinitiation complexes (20-22). These findings suggest that the core promoter and associated basal transcription factors are an important yet unexplored component in the regulation of gene expression, controlling differentiation transitions.

In this study we aimed to explore, in an *in vivo* model of development, the role the core promoter plays in defining transcriptional output in cells undergoing differentiation coupled changes in cell cycle dynamics, through both promoter-level regulatory and behavioural changes. The zebrafish Fucci transgenic system was used to ask how cell cycle dynamics affect promoter usage during differentiation stages. We have segregated cells as they go through differentiation coupled changes in cell cycle dynamics in somitogenesis stage embryos, which are characterised by marked spatial separation of differentiating cells with distinct cell cycle dynamics (18). CAGE 5’ end transcript analysis was used to interrogate the promoter features of these cell populations, in order to identify changes in the usage of transcriptional regulatory machinery. This investigation identified that the Fucci system can successfully be used to segregate cells on the basis of cell cycle stage and cycling behaviour, as well as differentiation status, when this is marked by changes in cell cycle dynamics. Interrogation of promoter behaviour between populations revealed a distinct sharpening (condensation of TSS usage to a narrower region of the promoter) in promoter usage, in genes upregulated in slowly cycling, differentiating tissues. This event was associated with enhanced utilization of the TATA-box, in addition to Sp1 binding sites, in particular the GC-box. A concurrent enhancement in CCAAT box utilization in genes upregulated in rapidly cycling cells, was also observed, in genes with a broad range of TSS utilization in particular. A greater utilisation of TATA-like and W-box motifs was also identified in rapidly cycling cells. A similar pattern of regulatory motif utilisation was observed in genes transitioning in their TSS distribution and with differentially used alternative promoters. This demonstrates a switch in core promoter associated transcriptional regulatory machinery utilization, leading to changes in promoter behaviour, as cells go through differentiation coupled changes in cell cycle dynamics. This investigation explores, for the first time, the regulation of tissue-lineage-specification on the promoter-level, in a whole organism, *in vivo* context.

## Materials and Methods

### Zebrafish husbandry and embryo generation

All animal husbandry and associated procedures were approved by the British Home Office (Licence number: P51AB7F76). Zebrafish embryos were obtained by sibling crosses from adult Fucci [(Tg(EF1α:mKO2-zCdt1(1/190))^rw0405b^ / Tg(EF1α:mAG-zGem(1/100))^rw0410h^] (18) fish housed in the University of Birmingham fish facility. Zebrafish were and bred and embryos raised and staged following standard protocols (14, 23).

### Light sheet imaging and image processing

Zebrafish embryos were decorionated using 1 mg/ml Protease from *Streptomyces griseus* (Sigma). Embryos were screened for fluorescence using a fluorescent microscope (Zeiss axio zoom V16), and mounted in a column of 1% low melting point agarose (Sigma) for light sheet imaging. Zebrafish embryos were imaged using dual side illumination, with maximum intensity projections (MIPs) processed images generated using the Zen Black imaging software (Zeiss).

### Cell sorting and RNA preparation

Zebrafish were decorionated and selected as previously described at the 14-somite stage. The embryos were dissociated into a single cell suspension using enzyme-free cell dissociation buffer, PBS based (Gibco). Dissociated cells were pelleted and resuspended in Hanks balanced salt solution without calcium chloride or magnesium sulphate (Sigma), for sorting. Cells were fluorescence-associated cell sorted (FACS) into populations displaying red and green fluorescence. Validation of correct sorting was performed by propidium iodide DNA content analysis following manufacturer’s conditions (Invitrogen). RNA was extracted from isolated cells using the miRNeasy kit (Qiagen). RNA quality was analysed by capillary electrophoresis (Bioanalyzer 2100, Agilent). All samples had an RNA integrity number (RIN) >9.

### CAGE library preparation and sequencing

NanoCAGE libraries were generated following a protocol described in (24). Pooled libraries, representing FACS sorted populations in triplicate were sequenced with the Illumina HiSeq 2500 50 cycles single-read run operation program, following manufacturer’s protocol.

### CAGE mapping and CTSS calling

The zebrafish genome assembly (Zv9) was downloaded from UCSC Genome Browser (25). Reads were trimmed (15 bp from 5’end) to remove the linker and UMI region. Reads were mapped using Bowtie (26), allowing a maximum of two mismatches and only uniquely mapping tags with MAPQ of 20. R/Bioconductor package CAGEr was used to remove the additional G nucleotide due to the CAGE protocol where it did not map to the genome (27). All unique 5’ ends of reads were defined as CAGE defined TSS’s (CTSS) and reads were counted at each CTSS per sample. These raw read counts were subsequently normalized based on a power-law distribution based on 10^6^ reads (28) and defined as normalized tags per million (tpm). After quality control per sample, a high level of inter-replicate correlation was observed and the biological replicates were merged for downstream analyses. One library of a biological replicate of 3 was excluded in the S/G2/M-phase merger, based on low complexity of the library.

### Calling transcriptional clusters

CTSS that were supported by at least 0.5 tpm in one of the samples were clustered based on a maximum allowed distance of 20 bp between two neighbouring CTSS. These transcriptional clusters (TCs) were then trimmed on the edges to obtain more robust boundaries of TCs by obtaining the positions of the 10th and 90th percentiles of expression per TC. Only TCs with higher than 5 tpm expression were considered. Finally, TCs across all three samples (G1, S/G2/M, total) were aggregated if within 100 bp of each other to form consensus clusters (CC) for downstream analyses.

### Annotation

The CCs were annotated to the nearest reported TSS from Ensembl (danRer7) using the R/Bioconductor package ChIPseeker (29). We selected only CCs that mapped within 1 kb upstream of the reported TSS as well as CCs mapping to 5’ UTR.

### Differential gene expression analysis

The raw read counts were extracted for the CCs across triplicates described earlier and collapsed into total count per CC. DESeq2 R/Bioconductor package was used to define differential expression and the threshold of differential expression was set at adjusted p-value of < 0.05. These results were cross referenced to the CC information of the merged samples. In cases of more than one CC mapping to the region, the CC with the highest expression was chosen to represent the region.

### Gene ontology

CCs were annotated with entrez gene IDs and analysed with GOstats R/Bioconductor package for overrepresentation of GO terms for biological processes. The up and down regulated genes were tested separately against all genes expressed amongst the samples.

### Tissue-specificity and cell cycle enrichment

The top 600 ranked human genes were downloaded from the Cyclebase database version 3.0. For each category of differential expression, the human orthologs (one to one & one to many) of zebrafish genes were determined using “mar2017.archive.ensembl.org” archive of hg19. The peak time phenotype was determined by Cyclebase. Enrichment of cell cycle genes was determined with a permutation test (n = 10,000). Tau and entropy scores for tissue-specificity were determined by RNA-seq expression of 8 adult zebrafish tissues from DanioCode series 391: brain, gill, heart, intestine, kidney, liver, muscle, and spleen. Analysis of the relative representation of tissue-specific terms, was performed by cross-referencing differentially expressed genes with mRNA *in situ* expression data extracted from the ZFIN database, for wild-type zygote to 20 days post fertilisation larval stages (https://zfin.org/downloads/wildtype-expression_fish.txt). This data was used to generate an anatomical specificity score that for each gene defines the pattern of spatial expression across development (Vucenovic and Lenhard, unpublished data). Data was then used to compare restrictedness of spatial localisation of gene expression for two groups of genes, stratified across developmental time.

### Core promoter motif enrichment analysis

Position weight matrices (PWMs) for TATA-box, CCAAT-box, GC-box, Sp1, INR, and YY1 were obtained from converting frequency matrices from JASPAR (7th release; 2018). Each CC was centred on the most expressed CTSS (the dominant TSS) and each sequence was scanned from 120 bp up and 50 bp downstream. A hit was reported if the scanned region contained a sequence with a 90% match to the PWN. For each group of differentially expressed genes occurrence was counted and compared to the non-significant set of CCs per sample. Significance was assessed using Fishers’ exact test. Obtained p-values were considered if < 0.01. The log2 odds ratios were visualised as a heatmap. Next, all possible variations of the pentamer using only T and A were generated for information on canonical TATA-box (TATAA) and other variants of the pentamer. Again, each sequence 20 to 40 bp upstream of dominant TSS was scanned for an exact match and counted for each occurrence. For each of the 2 groups, occurrences were counted and compared to the non-significant set of CCs per sample. Significance was assessed using Fishers’ exact test. Obtained p-values were considered if < 0.01. The results are visualized as the relative occurrence per group for the 11 most different pentamers between groups.

### Promoter shape classification

Width of CCs (interquantile-width, IQW) were defined as the distance between the positions of the 10th and 90th percentiles of expression per CC. Sharp promoters were characterised by a width < 10 bp, peaked broad promoters as >= 10 bp with one CTSS expressing more than 60% of expression of entire CC, and broad promoters as the remaining set.

### Alternative promoter utilisation

The genomic location of 1384 twinned canonical and alternative promoters identified in (30), were extracted and intersected with the G1 and S/G2/M CAGE data generated for this paper. Where multiple alternative promoters are present, only the highest expressed one was selected. TPM values within these regions were calculated and the expression patterns of genes with a TPM>5 in either the canonical or alternative promoter region in both the G1 and S/G2/M populations identified. Genes with a 2-fold change in the expression of the alternative promoter normalised to canonical promoter expression, were selected for further investigation (n=79). These genes were segregated for the relative behaviours of alternative and canonical promoters as follows. *Canonical down* (expression of the canonical promoter in S/G2/M cells is >50% decreased vs. G1 cells [S/G2/M cano down] and Δ [difference in expression] S/G2/M vs. G1 for the alternative promoter is <25%), *Alternative down* (Δ cano <25%, Δ alt >50% [S/G2/M alt down]), *Canonical up* (Δ cano >50% [S/G2/M cano up], Δ alt <25%), Alternative up (Δ cano <25%, Δ alt >50% [S/G2/M alt up]). Situations falling outside of these criteria were discarded (n=9).

## Results

### Segregation of embryonic cells with distinct cell cycle dynamics

The FUCCI transgenic system, differentially marks cells in G1 and S/G2/M phases of the cell cycle (Figure 1A) and can therefore be used to separate rapidly and slowly cycling cells *in vivo*, by virtue of the cell cycle stage they primarily inhabit. Sugiyama et al., 2009 (18) demonstrated a switch in the ratio G1 vs. S/G2/M marked cells in zebrafish undergoing somitogenesis, from primarily S/G2/M in early stages, to primarily G1 in later stages, associated with cell differentiation. In order to further investigate this observation, longitudinal assessment of cell proliferation rate during zebrafish embryo development, using this system, was performed and revealed a spatial and tissue-specific separation in cell cycling behaviour across post-gastrulation embryos (Figure 1B, Supplementary figure 1A, Supplementary movie 1), with cells, primarily in the developing somites, slowing their cycling, displayed by an elongated period in G1-phase and therefore an accumulation of red fluorescent signal in the somites. In contrast, green (S/G2/M) cells mark primarily neuroectoderm derived lineages, such as optic cup, neural tube and notably clearly identifiable cells in the notochord and circulating cells over the yolk ball (Figure 1B, Supplementary figure 1A, Supplementary movie 1). 14-somite embryos show the clearest spatial and tissue-specific segregation of cells on the basis of cell cycling dynamics (Figure 1B, Supplementary movie 2 and 3). In order to investigate the role of promoter associated transcriptional regulatory machinery in defining this transition, the 14-somite stage was selected for further investigation.

**Figure 1.**
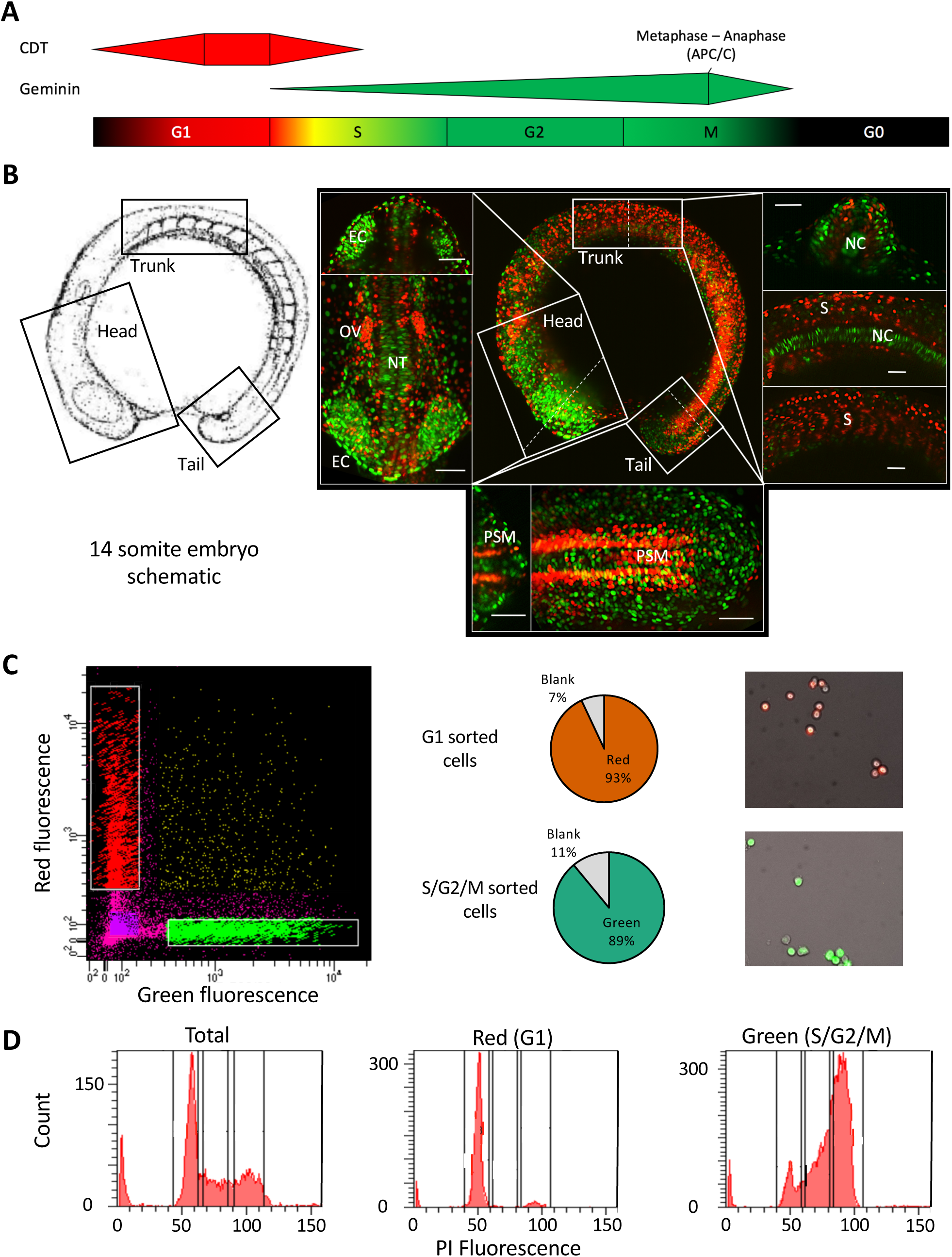
FACS mediated separation of cells by cell cycle stage in the developing embryo. [A] Schematic of fluorescence from the FUCCI system through the cell cycle with colours indicating phases marked by the fluorescent reporter genes. [B] Left panel; a schematic of the 14 somite embryos, reproduced with permission from (14). Right panel; center: max projection fluorescent image of a 14 somite FUCCI transgenic embryo, showing the distribution of rapidly (green) and slowly cycling (red) cells. Surrounding panels: higher magnification views of the head, trunk and tail regions with alternate views shown in the top/right sections and cross-sections (denoted by the dashed lines) shown in the bottom/ left sections of each surrounding panel. Scale bar = 50 uM. Abbreviations, Otic vesicle (OV), Eye cup (EC), Notochord (NC), Somites (S), Neural tube (NT), Pre-somitic mesoderm (PSM). [C] FACS sorting of FUCCI embryos, from left to right, FACS plot showing gating, pie charts showing cell selection efficiency and representative fluorescent images of isolated cells. [D] Propidium Iodide DNA content analysis of isolated cells. FACS traces show a shift in isolated cell DNA content from a level in accordance with primarily Gap phase 1 cells (G1) (gated as P3) in the red population to primarily S and G2 phase cells (gated in P4 and 5) in the green population, emanating from a mixed total population. Proportions are shown in Supplementary table 1.

To this end, 14-somite FUCCI embryos were dissociated to a single cell suspension and segregated by fluorescence associated cell sorting (FACS) into cells in G1 (red) and S/G2/M (green) (Supplementary figure 1B), with correct sorting confirmed by fluorescent imaging of the cells (Figure 1C). In order to validate that this process successfully segregates cells on the basis of cell cycle stage, segregated cells were subjected to DNA content analysis (Figure 1D). This analysis showed a marked enrichment for diploid (2N / G1) cells in the red, G1 segregated population over the total background population (81 vs. 39%) and enrichment for both 2-4N (S-phase) and tetraploid (4N / G2/M-phase) in the green, S/G2/M segregated population over total (27 vs. 17% and 46 vs 22% respectively) (Figure 1D, Supplementary table 1). This result confirms the successful segregation of cells on the basis of cell cycle stage using the FUCCI-transgenic system.

### Global transcription initiation patterns at known promoters

Next, we asked about the state of the mRNA transcriptome in fast and slow dividing cells of the embryo, with particular focus on the mRNA 5’ end in order to identify features of promoter utilisation. To achieve this goal, we chose a small cell number optimised protocol for detection of mRNA 5’ ends (nanoCAGE (24)), which reports steady state mRNAs quantitatively, and simultaneously informs about TSS usage and core promoter architecture (31). NanoCAGE was performed on three biological replicates of G1 and S/G2/M-phase segregated cells from the 14-somite stage zebrafish embryo, together with unsegregated cells (Total). CAGE reads were mapped to the zebrafish genome assembly (Zv9) and CTSSs assigned with a high level of inter-replicate correlation observed, with the exception of the S/G2/M (green) replicate 2, which was consequently excluded from further analysis (Supplementary figure 2, Supplementary table 2). Based on this the biological replicates were merged for downstream analysis.

In order to validate that this approach successfully identifies the transcription start sites, the distribution of mapped CTSSs was compared with previously published CAGE data (generated using the tagging-CAGE version of the protocol (32), from the 14 somite stage (30). Identified TSSs from each nanoCAGE sample, along with the tagging-CAGE, were grouped into consensus clusters (CCs) between samples and the distribution of TSSs (interquantile width [IQW]) within well expressed clusters (TPM≥5) compared. This analysis revealed very similar interquantile width distributions between the nanoCAGE data and previously published data (30) (Figure 2A, Supplementary figure 2B, Supplementary table 3). This is additionally exemplified by the very similar TSS distribution between samples of *si:ch211-113a14.29* (an orthologue of human histone 2B) and *mmp30* (matrix metallopeptidase 30) (Figure 2B). The determination of TSS distribution interquantile width is an established method for determining promoter shape, an important comparator of TSS utilisation (3-6).

**Figure 2.**
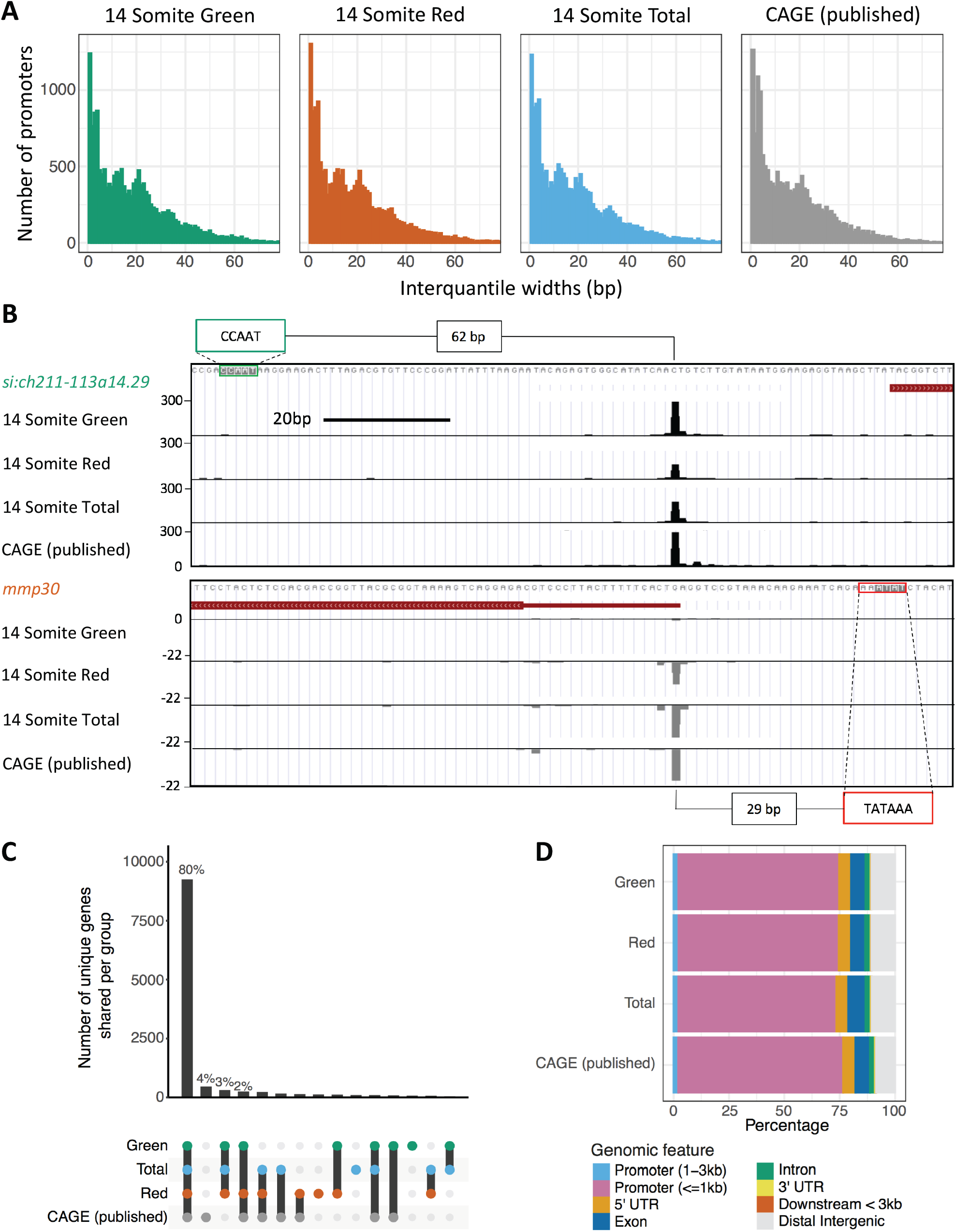
Overview of CAGE samples. [A] Histogram plots of the interquantile width of each Tag Cluster (TC). [B] UCSC genome browser view of *si:ch211-113a14.29* (an orthologue of human histone 2B) and *mmp30* (*matrix metallopeptidase 30*) showing consistent TSS distribution between nanoCAGE and published traditional CAGE. The position of putative regulatory motifs driving transcription are marked with distance from start of motif to dominant TSS position quantified. [C] A visualization of the overlap of TCs between the samples. The number of unique genes shared per group is shown in the lower half of the graph. [D] The number of TCs per sample mapping to genomic features. “Exon”, “5’ UTR”, “3’ UTR” and “intron” locations were extracted from the DanRer7 genomic build. “Promoter (<=1kb)” = window 0-1kb upstream of the gene start site, “Promoter (1-3kb)” = 1-3kb region upstream of gene start, “Downstream <3kb” = window 0-3kb downstream of the gene end annotated in the DanRer7 genomic build, and “Distal intergenic” = all regions not covered in other classifications.

A high degree of overlap was additionally observed for cluster position between nanoCAGE and previously published data, with 80% the same genes represented in all samples (Figure 2C, Supplementary tables 4&5). Identified CTSSs were additionally mapped to genomic features (Figure 2D, Supplementary figure 2C). The majority of CCs should fall within the core promoter region, <1kb upstream of the annotated 5’ end of genes, as this is known to be the major site of transcriptional initiation and accordingly in this analysis ∼70% of TSSs mapped to this region (Supplementary table 2), a similar level to previously published CAGE (Figure 2D). Taken together, these results indicate efficient isolation of gene promoter activities in cycling cells of differentiating embryos.

### Transcriptomes of G1 and S/G2/M cells reflect differential cell cycle and tissue-specific identities

CAGE is comparable to RNA-seq as a robust tool for quantitative transcriptomic analysis (30, 31, 33). Therefore, in order to profile the identity of cells segregated by cell cycle dynamics, and to identify differentially regulated genes, the expression of the promoter-associated consensus clusters was compared between samples. 12,865 consensus clusters were found to be shared between G1 (red) and S/G2/M (green), and clusters with a significant (P[adj]<0.05) change in expression between the populations identified (n=190 [up regulated in G1], n=138 [up regulated in S/G2/M]) (Figure 3A, Supplementary Table 6). This result indicates a large degree of overlap in the transcriptomes of the two populations, while differential regulation of a subset of transcripts opens the way to address their promoter regulation.

**Figure 3.**
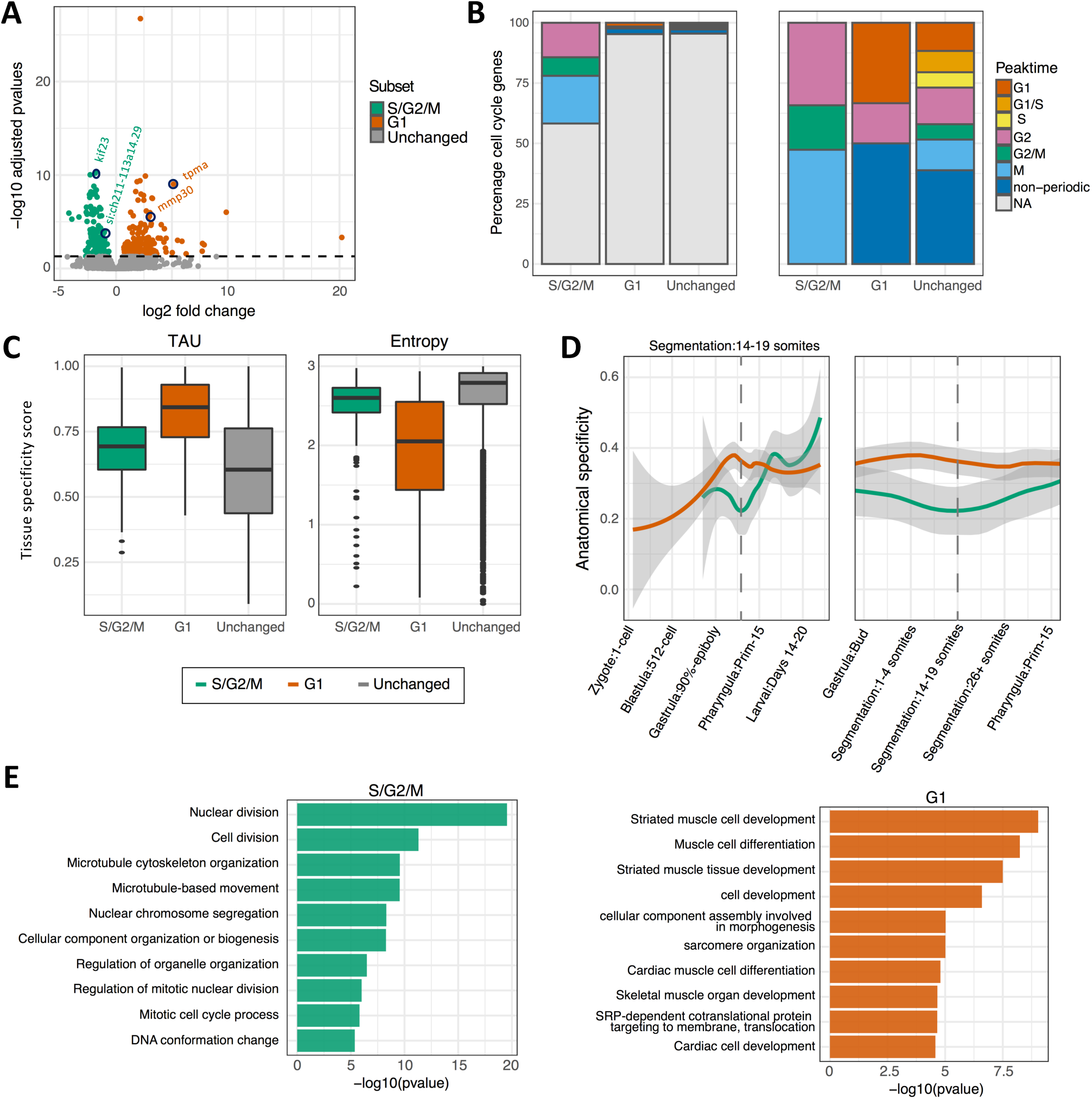
Classification of genes differentially expressed between G1 and S/G2/M segregated cells. [A] Volcano plot of all consensus clusters (CCs) in known promoter regions, coloured by significance. Location of genes shown in figure 2B as well as the identity of top significant differentially expressed genes between populations highlighted by circles and gene names. [B] Bar plot of the percentage of promoters overlapping a human annotated cell cycle periodic gene from Cyclebase for each differentially expressed group and the CCs unchanged between the two groups. Left panel, full sample groups (n= 138, 190 and 8406, S/G2/M, G1 and unchanged respectively). Right panel, cell cycle periodic genes (n= 43, 6 and 246, S/G2/M, G1 and unchanged respectively). [C] Box plot of tissue specificity scores, based on adult zebrafish tissue for genes upregulated in G1 and S/G2/M and unchanged. [D] Anatomical specificity scores, based on expression in embryonic zebrafish tissues, for genes upregulated in G1 and S/G2/M. Left panel, anatomical specificity scores across development with key stages marked on x-axis. Right panel, anatomical specificity scores around stage at which the samples were collected (14 somite). Grey shading marks standard error, dashed line marks 14-19 somite stage. [E] Gene ontology of biological processes with their corresponding p-values.

Fluorescence microscopy analysis of cell cycle dynamics in embryo development, revealed significant lineage-specific segregation of cells on the basis of their cell cycle dynamics (Figure 1B). Notably, the Fucci system indicates highly dynamic variation in the distribution of cell populations segregating into fast and slow dividing groups and domains in the embryo (Supplementary movie 2 and 3). Therefore, differentially expressed genes in cells segregated by cell cycle stage, in this context, may represent both cell cycle and lineage-specification related differences. In order to determine the contribution of each of these elements to the populations of differentially expressed genes, they were cross-referenced with a databases of cell cycle periodic genes (CycleBase) (Figure 3B) and tissue-lineage-specific genes, as determined by expression patterns in both adult and embryonic tissues (Figures 3C and D respectively).

Analysis of differentially expressed genes revealed a significant enrichment for cell cycle periodically expressed genes upregulated in the S/G2/M (green) population (46%, p<0.0001) with a peak of expression between G2 and M phase of the cell cycle (Figure 3B, Supplementary table 6). Conversely only 3.2% of genes upregulated in the G1 (red) population were cell cycle periodic. A significant majority of these however had peak expression in G1. This is in contrast to the group of genes with unchanged expression, where the periodicity of genes is more evenly distributed between cell cycle stages (Figure 3B).

We hypothesized that differential expression in cells with distinct cell cycle dynamics is a result, at least in part, to distinct cell cycle behaviour of cell-lineages, which are distinct in nature as well as represent varying levels of cell differentiation state. To test this, we asked about the contribution of lineage differentiation to selective expression of genes. Analysis of the tissue-lineage-specificity of differentially expressed genes revealed a clear enrichment in the tissue-lineage-specificity of genes upregulated in the G1 (red) population over the other populations (demonstrated by a higher average Tau and lower average entropy score) (Figure 3C). Analysis of the anatomical specificity of these gene lists across embryo development, revealed this enrichment to be highly specific to the segmentation stage of development, from which the samples were collected (Figure 3D). These results suggest that a significant contributing factor, leading to differential expression between the populations, is cellular replication on the part of the S/G2/M (green) population and tissue-specification on the part of the G1 (red) population. This observation is supported by the gene ontology of the differentially expressed genes, revealing in the S/G2/M (green) population, a clear enrichment for genes involved in DNA and chromatin processing, key for rapidly cycling cells, and in the G1 (red) population, enrichment for muscle development associated gene expression (Figure 3E).

In order to further dissect the contribution of different cell types and developing lineages to the slowly vs rapidly cycling (G1 and S/G2/M) populations, differentially expressed gene sets were cross-referenced with the ZFIN database of gene with known tissue and spatial specific expression at the 14-19 somite stage (Supplementary figure 3). The majority of genes did not match tissue-specific terms (72% [S/G2/M], 61% [G1] and 81% [Unchanged]) representing the fact that a minority of genes are tissue-specific and many tissues have not specified at the stage. This data does demonstrate however that the differentially expressed genes between the G1 and S/G2/M populations contain a disproportionate number of tissue-specific terms, particularly in the G1 population, in agreement with previous findings. Analysis of the relative contribution of genes with tissue-specific expression revealed a strikingly divergent expression pattern between populations (Supplementary figure 3), with somite and muscle specific terms highly enriched in the G1 population in agreement with gene ontology analysis and previously discussed fluorescence imaging (Figure 1B). Representation of terms related to the viscera, peripheral tissues (such as the periderm) and extra-embryonic tissues was also enriched in the G1 population, representing a population of cells starting to specify tissues within the embryo. In agreement with gene ontology analysis (Figure 3E), representation of proliferative and pluripotent cell types (such as the germ layers and proliferative region) was greatly enriched in the S/G2/M population. Interestingly, fluorescence imaging analysis (Figure 1B), suggest spatial distribution bias in the S/G2/M population compared to the G1 population, thereby S/G2/M phase green cells are particularly enriched in the eye cup and neural tissues. This apparent tissue bias is not borne out on the transcriptional level however, with neural and sensory tissue terms fairly evenly represented between populations (Supplementary figure 3). A closer inspection of the tissue distribution of the red green cells suggest that neural lineages share both fast and slowly dividing cells. Global transcriptomic analysis will reveal most enriched tissues, but will not reflect tissue-specificity of cell cycle regulation. Nevertheless, this analysis shows that the dynamics of cell cycle regulation follows certain trends, which manifests as an enrichment for the transition to a tissue-specific expression profile, marked by cell cycle dynamic changes, as seen in the slowly cycling, G1 population.

### Genes differentially expressed between G1 and S/G2/M cells utilise different core-promoter classes and regulatory elements

As revealed by the analyses detailed above, a clear transition in cell population identities is occurring during somitogenesis in these embryos, marked by a dramatic phenotypic change (speed of cell cycle) and a divergence in transcriptional output. In order to determine whether differential gene expression, associated with cell cycle dynamics, is also associated with changes in core promoter regulatory element distribution, the frequency of known, regulatory motifs 120 bp upstream and 50 bp downstream of the dominant TSSs of each promoter was determined. Both groups of differentially expressed genes showed marked changes in motif utilisation compared to genes with unchanged expression (Figure 4A, Supplementary table 7). In the S/G2/M population there is a statistically significant enrichment for the NF-Y factor associated CCAAT-box upstream of the TSS (p-value = 4.22 × 10^−12^). In the G1 population there is also a strong enrichment for a canonical TATA-box (p-value = 2.04 × 10^-7^), enrichment for Sp1 and GC-box (p-value = 0.002), and a depletion for a YY1 motif (p-value = 0.002). Strikingly, CCAAT box and TATA-box relative strength of enrichment are inverse for S/G2/M and G1 enriched genes, relative to those with unchanged expression (Figure 4A). Motif enrichment in each case was also specific, being positionally restrained relative to the TSS (Supplementary figure 4A). Examples of differentially expressed gene promoters, containing CCAAT/TATA-box motifs, are shown in (Figure 2B).

**Figure 4.**
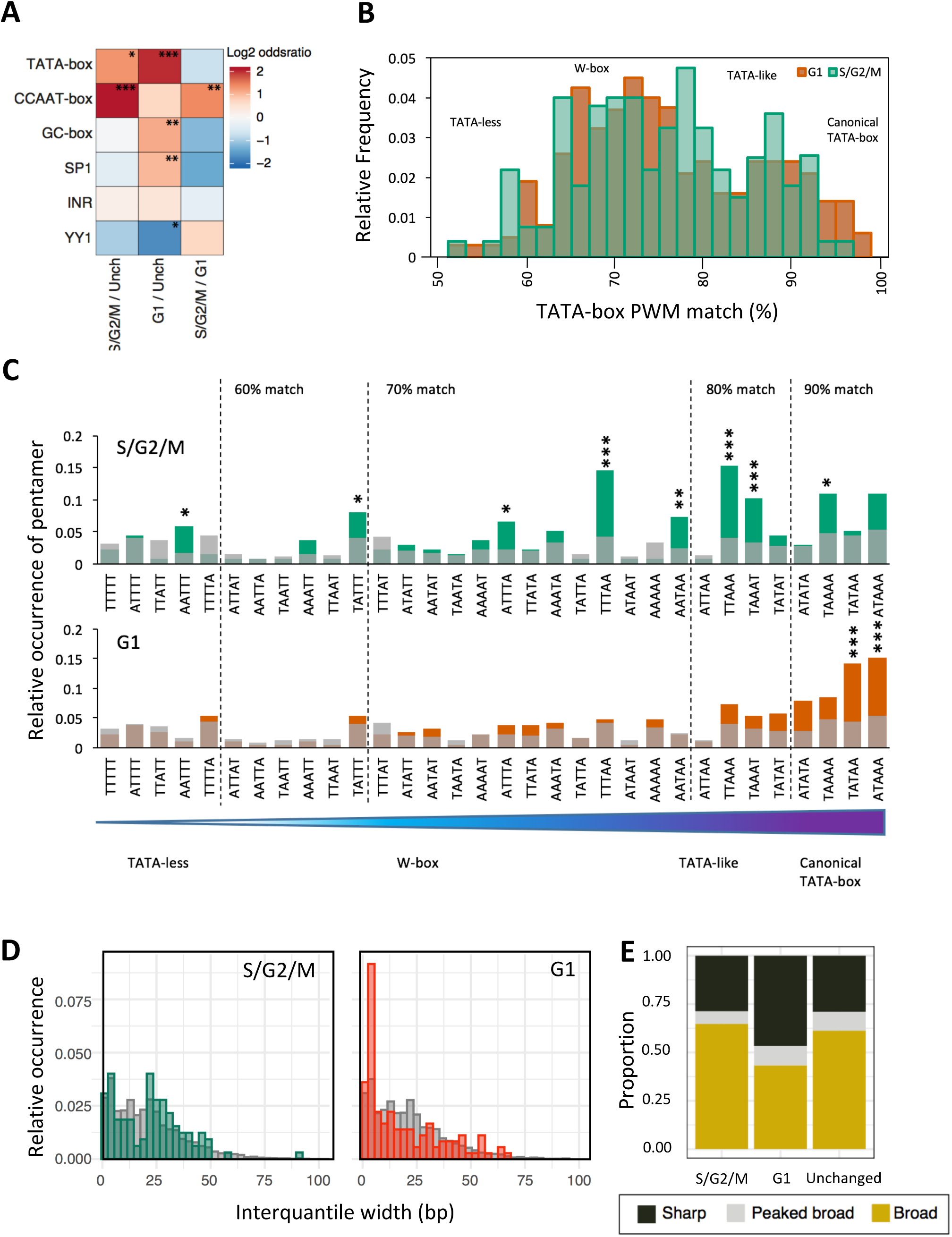
Core promoter architecture. [A] Heatmap visualizing the log2 odds ratio of occurrence of core promoter motifs between genes upregulated in G1 and S/G2/M and unchanged (*p<0.05, **p<0.01, ***p<0.001, Fisher’s exact test). [B] Distribution of position weight matrix (PWM) match (%) to TATA-box in the region –40 to –20 bp upstream of the dominant TSS in genes upregulated in S/G2/M and G1 populations. [C] A/T pentamer relative occurrence 20 to 40 bp upstream of TSS of genes upregulated in S/G2/M (green), G1 (red) and unchanged expression (grey). Ordering of pentamers is by best-fit match to the TATA PWM incremental thresholds shown. Pentamers significantly enriched in each group relative to pentamer occurrence in genes with unchanged expression (*p<0.05, **p<0.01, ***p<0.001, Fisher’s exact test). [D] Consensus cluster interquantile width in genes upregulated in S/G2/M (green) and G1 (red) and with unchanged expression (grey) visualised as a histogram. [E] Promoter shape distribution per group. Classifications; sharp (at least 90% of the expression from the promoter emanating from TSSs within 10bp of one another (IQW<10) and ≥60% of expression emanating from a single dominant TSS); Sharp with broad background (≥60% of expression emanating from a single dominant TSS, but IQW≥10) and Broad (<60% of expression emanating from a single dominant TSS and IQW≥10).

Beside altered utilisation of specific regulatory motifs, previous analyses of development linked changes in promoter utilisation, such as during zygotic genome activation (16), have found it to be associated with changes in the utilisation of the W-box, an A/T (WW) rich stretch with similar positional constraint as the TATA-box to the TSS. In order to investigate whether this motif is differentially used, the relative frequency of WW-dinucleotides and TATA-box (>90% PWM match), 120 bp upstream and 50 bp downstream of the dominant site of transcription, in differentially expressed genes was calculated (Supplementary figure 5). TATA-box frequency analysis was done alongside to differentiate TATA and W-box frequency, as a canonical TATA box will appear as a WW dinucleotide enrichment in this analysis. This analysis confirmed TATA-box enrichment in genes upregulated in G1 (∼30bp upstream of the TSS), but no enrichment in WW dinucleotide frequency between the G1 and S/G2/M differentially expressed gene sets (Supplementary figure 5). This suggests that TATA and A/T rich motifs (such as W-box and TATA-like) may be differentially used between G1 and S/G2/M populations. W-box and TATA-like motifs are distinct from the canonical TATA-box by virtue of a looser motif specificity (16, 34). In order to investigate their relative utilisation, the approximate location of the TATA-box (20-40bp upstream of the TSS) was analysed for TATA-box position weight matrix (PWM) match (Figure 4B, Supplementary figure 4B). This analysis revealed a shift from canonical TATA (>90% match) utilisation enriched in G1, to 75-90% match (previously identified as distinct TATA-like and W-box motifs (16, 34)) enriched in S/G2/M. TATA-like and W-box motifs constitute a highly diverse population in terms of sequence identity. Analysis of the relative occurrence of poly-W pentamers, 20-40bp upstream of the TSS, revealed that specific forms of TATA-like (TTAAA, TAAAT) and W-box (TTTAA, AATAA) are significantly enriched (p<0.001-0.01) in the promoters of genes upregulated in S/G2/M cells (Figure 4C, Supplementary table 8). Overall this suggests a change in the degree of utilization of canonical vs. non-canonical TATA regulatory elements in genes differentially expressed between G1 and S/G2/M populations.

Canonical TATA-box utilization is strongly associated with sharp promoters, where a single or condensed cluster of TSSs are used in the promoter (3-5) and is associated with high level of expression often associated with structural genes (6). In order to determine whether enhanced TATA utilization is associated with a difference in the shape of promoter utilization, the distribution of TSSs (interquantile width [IQW]) within clusters was determined and compared between samples (Figure 4D). Additionally, consensus clusters were classified into classes on the basis of the pattern of TSS utilisation (Figure 4E). Classifications were as follows; sharp (at least 90% of the expression from the promoter emanating from TSSs within 10 bp of one another (IQW<10) and ≥60% of expression emanating from a single dominant TSS); Peaked broad (≥60% of expression emanating from a single dominant TSS, but IQW≥10) and Broad (<60% of expression emanating from a single dominant TSS and IQW≥10) (3). This analysis identified a distinct enrichment in sharp promoter utilisation in G1 vs. S/G2/M and non-significantly differentially expressed genes (43.7% vs 29.0% and 29.2% respectively) (Figure 4E). In order to determine whether this relative sharpening of promoter utilisation in G1 differentially expressed genes was due to increased TATA utilisation, relative TATA motif frequency (>90% PWM match) was determined in a promoter proximal region (120 bp upstream and 50 bp downstream of the dominant site of transcription) in G1 vs S/G2/M differentially expressed genes, as previously described, but segregated by promoter shape (sharp, peaked broad and broad) (Supplementary figure 5). This analysis showed that indeed TATA-box utilisation is highly enriched in sharp promoters over other behaviours, interestingly however this is only true in the gene set upregulated in G1. In the S/G2/M upregulated gene set promoter shape was only weakly associated with TATA utilisation (Supplementary figure 5). WW-dinucleotide frequency analysis done in an identical manner also showed no association with promoter shape. Collectively these findings suggest that enhanced TATA utilisation and a greater proportion of sharp promoters are associated in genes upregulated in slowly cycling G1 cells (Figure 4D and E, Supplementary figure 5).

### Gene promoters with marked change in TSS distribution between populations display differential regulatory element utilisation

Analysis of genes differentially expressed between the G1 and S/G2/M population (Figures 3 and 4) show distinct differences in promoter behaviour (sharper promoter utilization in slowly cycling G1 cells) and the regulatory networks associated with their expression (often involving the differential usage of TATA or CCAAT boxes). As described in Figure 2A, the global distribution of promoter usage is unchanged between populations, however analysis of TSS distribution, in promoters highly expressed (TPM>10) in both populations, revealed a significant proportion of genes transitioning in promoter shape (sharp, peaked broad and broad) between populations (394/4774, 8.3%) (Figure 5A). This event is potentially representative of a transition in dominant promoter regulatory network (3-5). *Mitochondrial fission regulator 2* (*mtfr2*) for example, a gene associated with cell proliferation in human (35), displays a shape change transition between populations, broadening TSS distribution in the S/G2/M population vs. G1. This gene additionally contains a CCAAT box within its promoter, proximal to the shape change event (Figure 5B).

**Figure 5.**
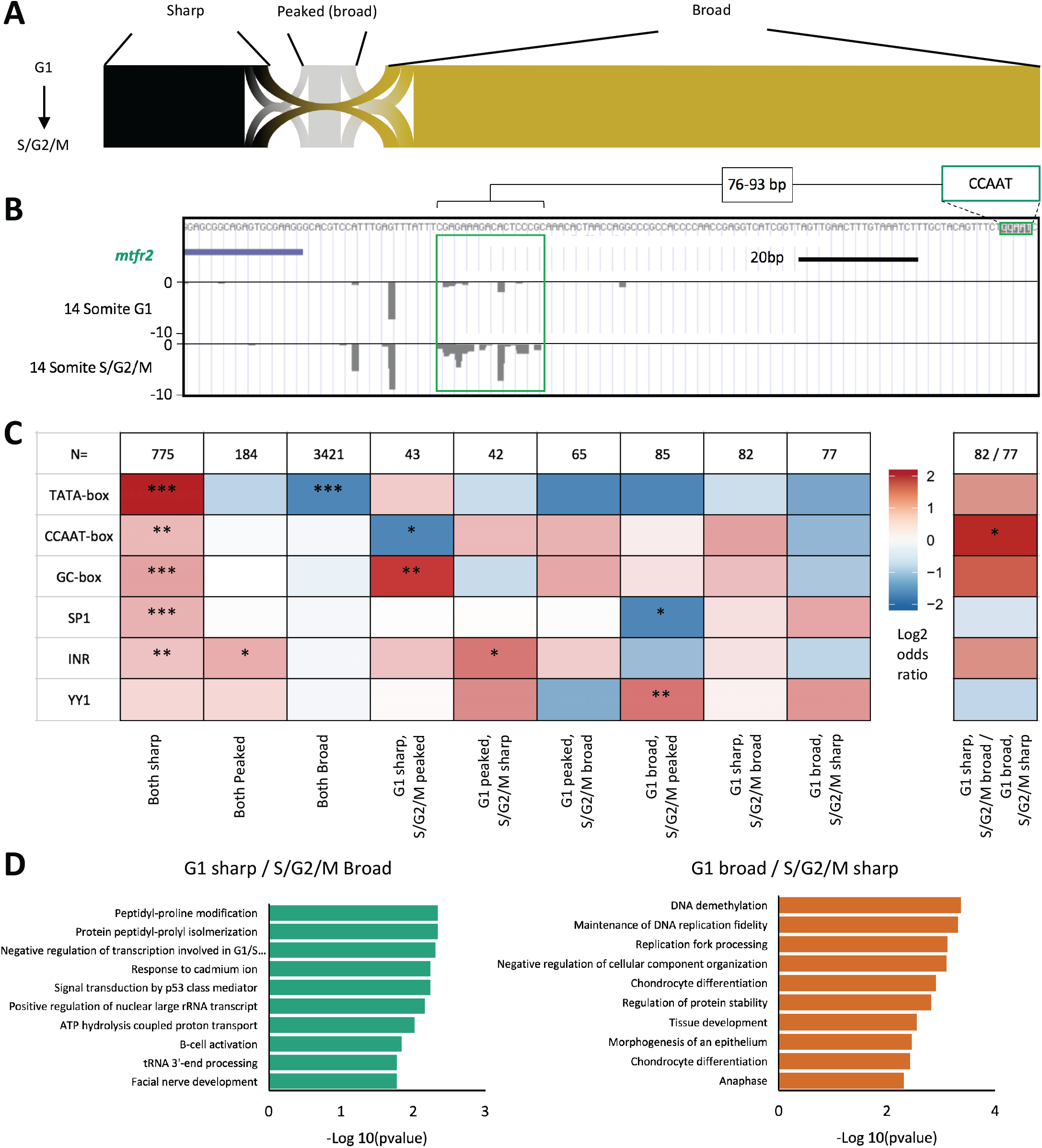
Core promoter shape transition. Promoter interquantile width (IQW) was measured and dominant TSSs assigned for consensus clusters (CCs) with at least 10 tpm expression in both the G1 and S/G2/M populations (n=4774). Promoters were segregated into 3 groups based on promoter TSS distribution (shape), sharp (IQW<10bp), peaked broad (IQW>10bp, dominant TSS >60% of CC expression) and broad (IQW>10bp, dominant TSS <60% of CC expression). [A] Sankey plot of promoter shape correspondence between G1 and S/G2/M. [B] UCSC genome browser view of the *mitochondrial fission regulator 2* (*mtfr2*) promoter with a shape change transition between populations, broadening TSS distribution in the S/G2/M population vs. G1. A proximal regulatory element (CCAAT box) is highlighted along with its spatial proximity to the main region of differential TSS utilisation in this promoter (green box). [C] Heatmap visualizing the log2 odds ratio of selected promoter motif occurrence for promoters transitioning in shape. Left heat map, occurrence is scored in each group vs. the rest of the dataset of shared CCs >10tpm expression (n=4774). Right heatmap, comparison is as shown. (*p<0.05, **p<0.01, ***p<0.001, Fisher’s exact test). [D] Gene ontology of biological processes of genes with promoters transitioning from sharp to broad between populations, with corresponding p-values shown.

This population of promoters show context dependent changes in their utilisation. They therefore could represent an important population defining alterations in the promoterome (and therefore the transcriptome) of cells undergoing differentiation coupled changes in cell cycle dynamics. In order to determine whether, like differentially expressed promoters, these transitioning promoters are marked by different regulatory machinery to each other, promoter proximal regulatory motif analysis, was performed, as before (Figure 5C). This analysis revealed, in line with previous studies (6-9), significant enrichment for regulatory motifs (particularly TATA) in genes that remained sharp in both populations and a depletion of TATA in genes where broad promoter utilisation is retained (Figure 5C). Of note YY1 motif occurrence is enriched in genes with a condensed TSS distribution in the S/G2/M population relative to G1 (G1 peaked / S/G2/M sharp, G1 broad / S/G2/M sharp, G1 broad / S/G2/M peaked) the latter significantly (n<0.01). Of greatest interest however are gene promoters where there is a significant shift in TSS distribution (from sharp to broad) between populations. Direct comparison of these populations (G1 sharp / S/G2/M broad vs. G1 broad / S/G2/M sharp) revealed a significant (p<0.05) enrichment for CCAAT box occurrence in the former population (Figure 5C), further supporting the association of the CCAAT box motif with differential promoter usage between populations. Gene ontology analysis of these broad-to-sharp shifting populations revealed a mixed set of terms with both proliferation and differentiation genes represented (Figure 5D). This suggests that transitions in the TSS distribution on the promoter has a subtler effect on transcriptional output, than on differentially expressed genes, however both processes are driven by the utilisation of similar regulatory motifs and argue for distinct activity of general transcription factor complexes on promoters in fast and slow dividing cells.

### Differential utilisation of alternative promoters between cell cycle dynamic divergent populations

A major source of divergence in the transcriptome of cell populations, particularly during differentiation, is through the use of alternative promoters (30). In order to investigate whether the regulatory differences, observed with differentially expressed and promoter shape transitioning genes, also impact on the utilisation of alternative promoters, the relative expression of previously identified alternative promoter containing genes in zebrafish (30), was determined between populations. TPM values were calculated for genomic regions identified to correspond to canonical and alternative promoters associated with the same genes (n=1384). Genes with significant expression (TPM>5 in either the canonical or alternative promoter region in both the G1 and S/G2/M populations) were selected for further analysis (n=231). In order to identify genes with differential utilisation of alternative promoters, alternative promoter TPM values were normalised to canonical (alternative promoter relative expression) and compared between populations (Figure 6A). This analysis identified 79 genes (34% of significantly expressed candidates) with a 2-fold change in the relative expression of the alternative promoter.

**Figure 6.**
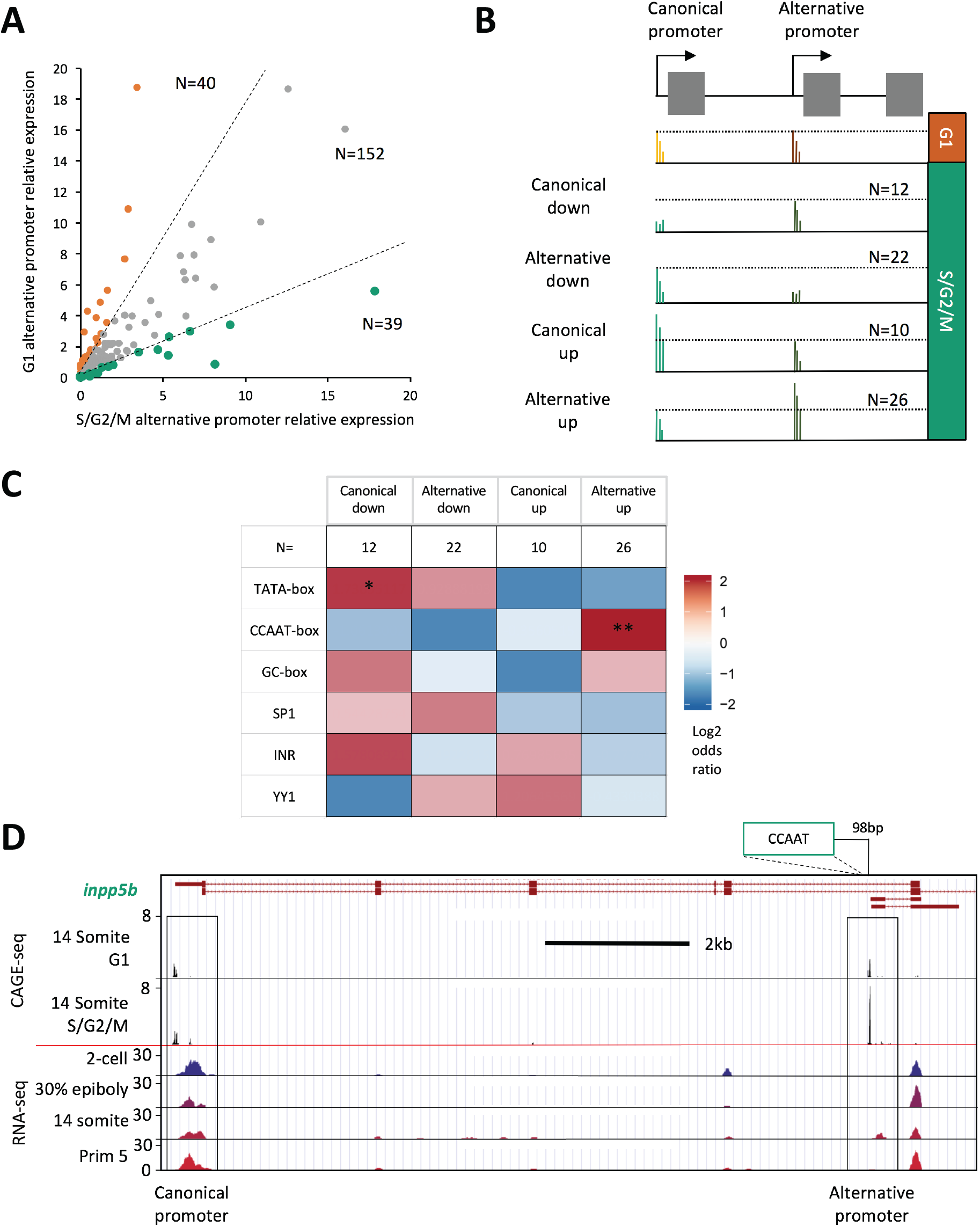
Alternative promoter usage. TPM values were determined for genomic regions identified in Nepal et al., 2013 (30) to correspond to canonical and alternative promoters, associated with the same genes (n=1384). Genes with significant expression (TPM>5 in either the canonical or alternative promoter region in both the G1 and S/G2/M populations) were taken for further analysis (n=231). [A] Plot of correlation of alternative promoter utilisation, normalised to canonical promoter expression, between G1 and S/G2/M populations. Dashed lines denote threshold of 2 fold change in normalised alternative promoter expression between the populations (n=79). These differentially utilised genes were segregated by whether a change in the expression of the canonical or alternative promoter, in either population, was responsible for the 2-fold change in the expression of the alternative promoter, normalised to canonical promoter expression. [B] Diagrammatic summary of group selection criteria. In brief, relative to the G1 population; *Canonical down*: S/G2/M alternative promoter expression unchanged, but canonical expression depleted (n=12); *Alternative down*: S/G2/M alternative promoter expression depleted, but canonical expression unchanged (n=22); *Canonical up*: S/G2/M alternative promoter expression unchanged, but canonical expression enhanced (n=10); *Alternative up*: S/G2/M alternative promoter expression enhanced, but canonical expression unchanged. Situations falling outside of these criteria were discarded (n=9). Full selection criteria shown in materials and methods. [C] Heatmap visualizing the log2 odds ratio of selected promoter motif occurrence for each group. Occurrence is scored in each group vs. occurrence in the rest of the data set (n=80). (*p<0.05, **p<0.01, ***p<0.001, Fisher’s exact test). [D] UCSC genome browser view of the *inositol polyphosphate-5-phosphatase B* (*inpp5b)* promoters with CAGE-seq tracks showing enhanced alternative promoter usage in the S/G2/M population vs G1, an example of a member of “Alternative up” group. The position of a CCAAT motif relative to the alternative promoter is shown. RNA-seq tracks, imported from the “Promoterome CAGE and nucleosome positioning” publicly available trackhub (URL: http://trackhub.genereg.net/promoterome/danRer7/index.html) (16, 30) show that this alternative promoter drives a somitogenesis specific transcript.

In order to determine which of these relative changes in expression were due to an upregulation in the canonical or alternative promoter in either the G1 or S/G2/M populations, these genes were subdivided into four groups as detailed in Figure 6B; They are as follows; relative to the G1 population; *Canonical down*: S/G2/M alternative promoter expression is unchanged, but canonical expression is depleted (n=12); *Alternative down*: S/G2/M alternative promoter expression is depleted, but canonical expression is unchanged (n=22); *Canonical up*: S/G2/M alternative promoter expression is unchanged, but canonical expression is enhanced (n=10); *Alternative up*: S/G2/M alternative promoter expression is enhanced, but canonical expression is unchanged. Situations falling outside of these criteria were discarded (n=9). Full selection criteria shown in materials and methods. In order to determine whether, like differentially expressed and shape changing promoters, these alternative promoter utilisation events are marked by differential regulatory machinery to each other, promoter proximal regulatory motif analysis, was performed, as before (Figure 6C). This analysis revealed a significant enrichment for TATA utilisation in events, where the canonical promoter is depleted in S/G2/M (P<0.05) and significant CCAAT utilisation enrichment where the alternative promoter is enriched in S/G2/M (P<0.01). An example of the latter is shown in Figure 6D. The utilisation of the canonical promoter of *inositol polyphosphate-5-phosphatase B* (*inpp5b)* is unchanged between population, however expression from an alternative, CCAAT box containing, promoter is enhanced in S/G2/M. Publicly available RNA-seq data from (16, 30) suggests that this promoter drives an alternative somitogenesis specific transcript from the *inpp5b* gene. This analysis further supports a role for differential TATA- and CCAAT-box core-promoter utilisation in defining disparate transcriptional output, between populations segregated by cell cycle dynamic behaviour.

## Discussion

The developing zebrafish embryo displays extensive transitioning in cell cycle dynamics associated with cell differentiation. Previous studies have described how this process is accompanied by changes in transcriptional output, but this study is the first to show the role the core-promoter plays in defining this output through local regulatory changes. In this investigation, cells from somatising FUCCI transgenic embryos were dissociated and segregated by cell cycle stage (G1 vs S/G2/M). Differential expression analysis on these populations revealed a striking separation in the identity of these cells. Beside differences in cell cycle stage, slowly cycling (G1) cells showed extensive specification in terms of gene expression to different tissues, associated with increased representation of tissue-specific regulatory motifs in their promoters. Rapidly cycling (S/G2/M) cells, on the other hand, were far less specified, with overwhelming representation of ubiquitously expressed genes, associated with DNA and chromatin processing required for rapid proliferation. This divergence in cellular identity was found to be concurrent with changes in promoter behaviour. Differentially expressed genes were sharper in the G1 population and associated with an increase in TATA-box utilisation, alongside GC-box and Sp1. In contrast the broader promoter usage in S/G2/M upregulated genes was accompanied by a far greater utilisation of CCAAT box general transcription factor binding sites and a lesser dependence on TATA. TATA-like and W-box motif frequency was found to be enriched in rapidly cycling cells however. Global analysis of changes in promoter associated TSS distribution (shape change) and usage of alternative promoters, revealed these again to be associated with divergent utilisation of TATA and CCAAT box TF binding sites.

Collectively this data suggests a strong divergence in the utilisation of general transcription factor binding sites, TATA and CCAAT as well as Sp1 binding sites, in particular the GC box, between cells undergoing differentiation coupled changes in cell cycle dynamics. This divergence impacts on the transcriptome and promoterome of these cells resulting in differential gene expression as well as differential canonical and alternative promoter utilisation.

Promoter associated regulatory elements have previously been associated with marking genes with cell cycle periodic expression. The promoter-associated cell cycle-dependent element (CDE) and the cell cycle genes homology region (CHR) have both been found to regulate genes with maximum expression in G2-M phase, through cell cycle stage dependent binding of transcription factors (reviewed in (36)). CCAAT-boxes have also been found to play a role, in association with these factors. Three CCAAT boxes, along with a single cell cycle gene homology region (CHR), were found to be major regulatory sites for the transcription of human cyclin B2, through NF-Y binding (37). Additionally CCAAT/enhancer binding protein β (C/EBPβ) was found to be a key driver of stromal cell progress through the cell cycle (38). This suggests that promoter based regulatory signalling is key to controlling gene expression cell cycle periodicity. This study extends this by displaying a role for CCAAT boxes in genes differentially regulated during differentiation coupled changes in cell cycle dynamics, associated with broad TSS distribution across the promoter.

This investigation shows that cells going through the striking changes in cell cycle dynamics, occurring embryo wide, during somitogenesis, have differential tissue-specification. This may suggest a marked transition in multiple tissue progenitors during this process. Rapidly cycling cells, show weak tissue-specificity despite their clear segregation to distinct tissue domains such as the eye cup notochord and brain tissues. Additionally, they show a transcriptional profile dominated by ubiquitously expressed DNA processing genes and cell cycle regulators (39). This population of genes has been found to be downregulated at the onset of organogenesis in mouse and shortly after gastrulation in drosophila (40), highlighting the modulation of this class of genes as a key marker of tissue-specification. Slowly cycling cells on the other hand show a far more tissue-specified transcriptional program, linked with a greater utilisation of the TATA-box associated gene regulatory machinery.

Single cell RNA-seq analysis of early stage embryo development (from high stage to 6 somite) does reveal an early specification point differentiating notochord, and brain tissues (both rapidly cycling) from somites and cardiac tissues (both slowly cycling) (41). This data supports the idea that the spatial segregation of cells based on cell cycle dynamics is associated with lineage-specification. As stated however, this analysis only extended to the 6-somite stage when the vast majority of cells are rapidly cycling (Supplementary figure 1A). In combination with the findings of this investigation, this data suggests that while lineages are specified early in development by small transcriptional changes (but with cell cycle and DNA processing genes predominant) and no discernable cell cycle dynamic changes, it is at the point of tissue differentiation that a major shift in transcriptional output occurs, associated with an elongation of the G1 phase. In the future, combining the approaches used in both investigations, possibly using newly developed single cell CAGE protocols (such as C1-CAGE) (42) will permit this process to be further explored.

Investigations into differentiation transition associated changes in gene regulatory programs, from terminally differentiated gamete, to pluripotent stem cell (16) and lineage defined multipotent stem cell, to terminally differentiated tissue cell (1, 19), have also found dramatic changes in gene regulatory program, associated with changes in the utilisation of TATA, TATA-like and W-box machinery (1, 16, 19). Terminally differentiated gametes reprogram to pluripotent cells in the early embryo, marked by W-box restricted programs being replaced with open CpG island associated promoter utilisation, restricted only by +1 nucleosome positioning, potentially priming pluripotent cells for a more diversified repertoire of behaviour as their lineage is specified (16). This process is also marked by a transition from rapid synchronous cell cycles, to slower asynchronous cycling, in zebrafish embryos. In this paper we show that cells then slow their cycling to differentiate and defining tissues as the body map starts to form, and this process is marked by an upregulation of TATA driven expression of tissue-specific genes and down regulation of ubiquitously expressed DNA and chromatin processing machinery. Studies, investigating transitions from lineage defined multipotent stem cells to terminally differentiated cells in both muscle (19) and liver development (1) have shown that this process is marked by wholesale depletion of RNA polymerase II regulatory cofactors, in particular TATA-associated factors. This is accompanied by a restriction to a small cohort of functional regulatory elements, in line with a limitation of the transcriptional repertoire, to be highly specialized and tissue-specific, in terminally differentiated cells. Collectively, this suggests that differentiation transitions in embryonic development are intimately associated with promoter level changes in the gene regulatory program, with the TATA-box a key component.

The TATA-box has previously been characterised as playing a crucial role in defining the tissue-specificity of associated genes, with precise TATA-TSS spacing a vital factor (4). Additionally, tissue-specific genes in *D. melanogaster* and mammalian systems are often associated with the presence of a promoter associated, spatially constrained TATA-boxes, alongside sharp transcription initiation, differentiating this class of promoter (type I) from non-TATA dependent housekeeping (type II) or developmentally regulated (type III) gene associated promoters (7, 43-45). This paper extends the tissue-specification/TATA-box association story by showing that an increase in TATA-box utilisation is one of the major defining factors in specifying differentiating cells at the earliest stages of tissue-lineage-specification.

Interestingly, in contrast to TATA-box utilisation, TATA-like and W-box motifs were found to be enriched proximal to the promoters of genes upregulated in rapidly cycling cells. TATA-like elements have been previously been associated with more dispersed TSS distribution on promoters, being associated with twin-TSS promoters (two dominant TSSs in the promoter) (4). Additionally, transcription mediated by TATA-like motifs had been found to predominantly regulated by the TFIID complex, rather than SAGA, in contrast to TATA-box regulation, in yeast (46). The regulatory distinction between TFIID and SAGA dominant genes has subsequently been challenged however, with SAGA reported to be a general cofactor required for virtually all RNA polymerase II transcription (47). This dependence on TFIID has be found to be subject to the presence of additional promoter regulatory elements including the upstream activating sequence (UAS) (48). The SAGA complex has not been identified in vertebrates, so how transferable this observation is to zebrafish development is uncertain, however it does demonstrate the TATA and TATA-like elements confer distinct regulatory identities to the promoters in which they reside. Significant and contrasting differences in the utilization of these motifs, identified in this study, in genes differentially expressed in fast and slowly cycling cells, suggests distinct promoter level regulatory programs activated between these populations.

Sp1, alongside its regulatory binding site, the GC-box, have also been associated with the regulation of tissue-specification, in particular through interaction with cell cycle regulated factors (49). Sp1 has been implicated in the transactivation of differentiation-regulated genes, underlying the switch from proliferation to differentiation, in squamous epithelium, through interactions with cell cycle regulators, retinoblastoma protein (pRB) and cyclin D1 (50). Alongside this, Sp1 has been identified as a key regulator of the G1 phase of the cell cycle in epithelial cells (51). This paper extends these observations by demonstrating the activity of Sp1 in the regulation of genes involved in the transition of cells from a proliferation to differentiation phenotype in a developing *in vivo* model.

This investigation demonstrates for the first time the promoter level regulatory changes occurring in cells undergoing the transition from proliferation to tissue-specification, *in vivo*. It displays how this process is coupled with transitions in cell cycle dynamics to a slower cycling rate where the G1-phase predominates, alongside transitions in promoter behaviour. This is marked by a sharpening of promoter utilisation, from rapidly to slowly cycling cells mediated by increased utilisation of TATA-box and Sp1/GC box regulatory units, at the expense of the CCAAT box. This transition in the promoterome is reflected in the transcriptome where the predominance of DNA and chromatin processing factors in rapidly cycling unspecified cells is replaced by increased expression of tissue-lineage specifying genes.

## Contributions

J.W. and F.M. conceived and coordinated the project. J.W. carried out all experiments and wet lab-based analyses. J.W. and L.R. analyzed sequencing data. J.W., L.R. and F.M. interpreted results with critical comments from N.C. and B.L. Anatomical specificity analysis was carried out by J.W. and D.V.

## Acknowledgements

The authors are grateful to Fabio D’Orazio and Sepand Rastegar for critical comments on the manuscript. This work was supported by the Wellcome Trust Investigator grant awarded to FM and BL.

---

**Supplementary figure 1.**
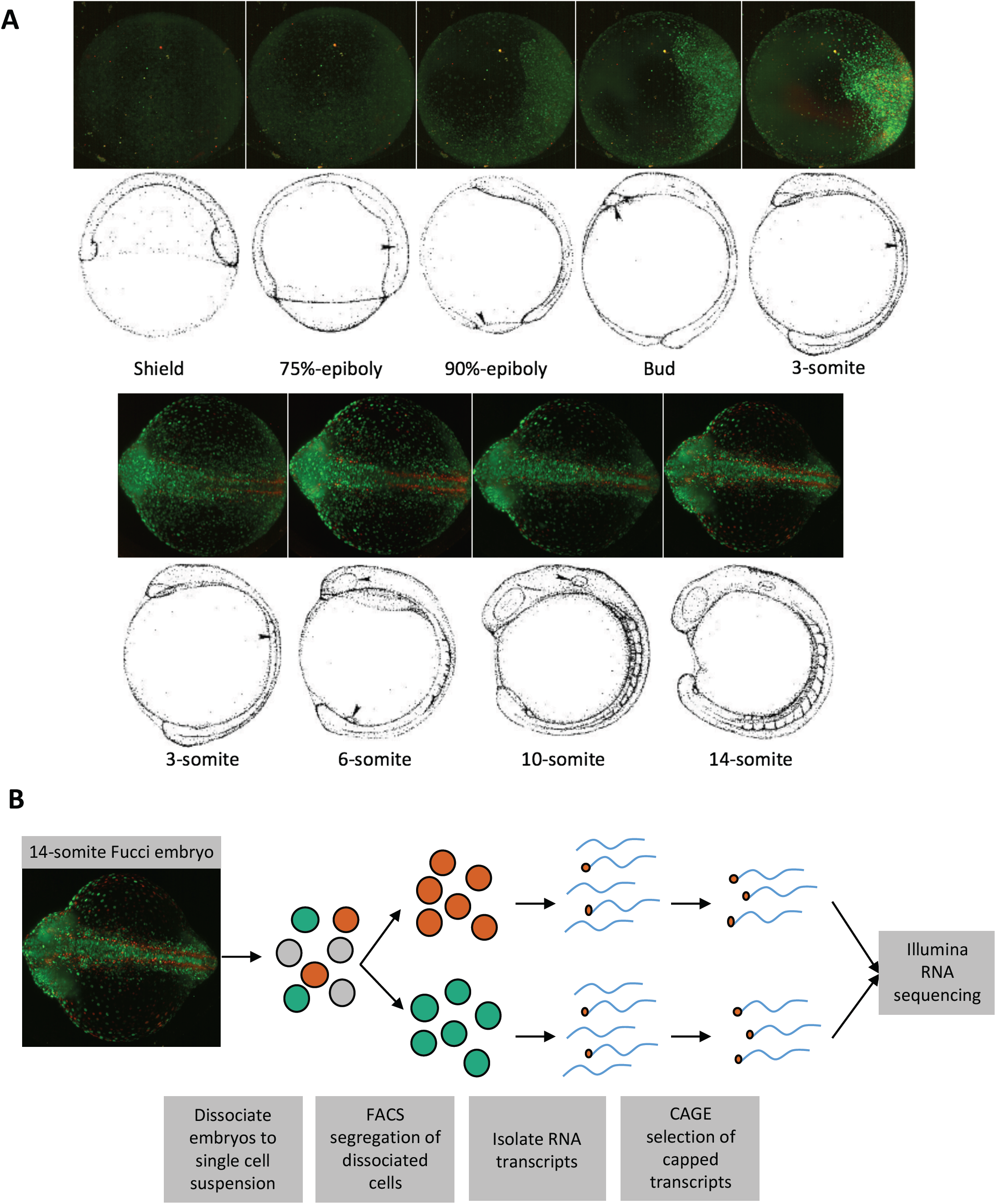
[A] Longitudinal analysis of FUCCI florescence in post-gastrulation embryos. Representative fluorescence images of FUCCI embryos from Shield to 14 somite stage. Drawings of embryo stage reproduced with permission from (14) [B] Flow diagram of experimental procedure for cycle-dynamics dependent segregation of cells and selection of capped transcripts for Illumina RNA sequencing and analysis of the promoterome.

**Supplementary figure 2.**
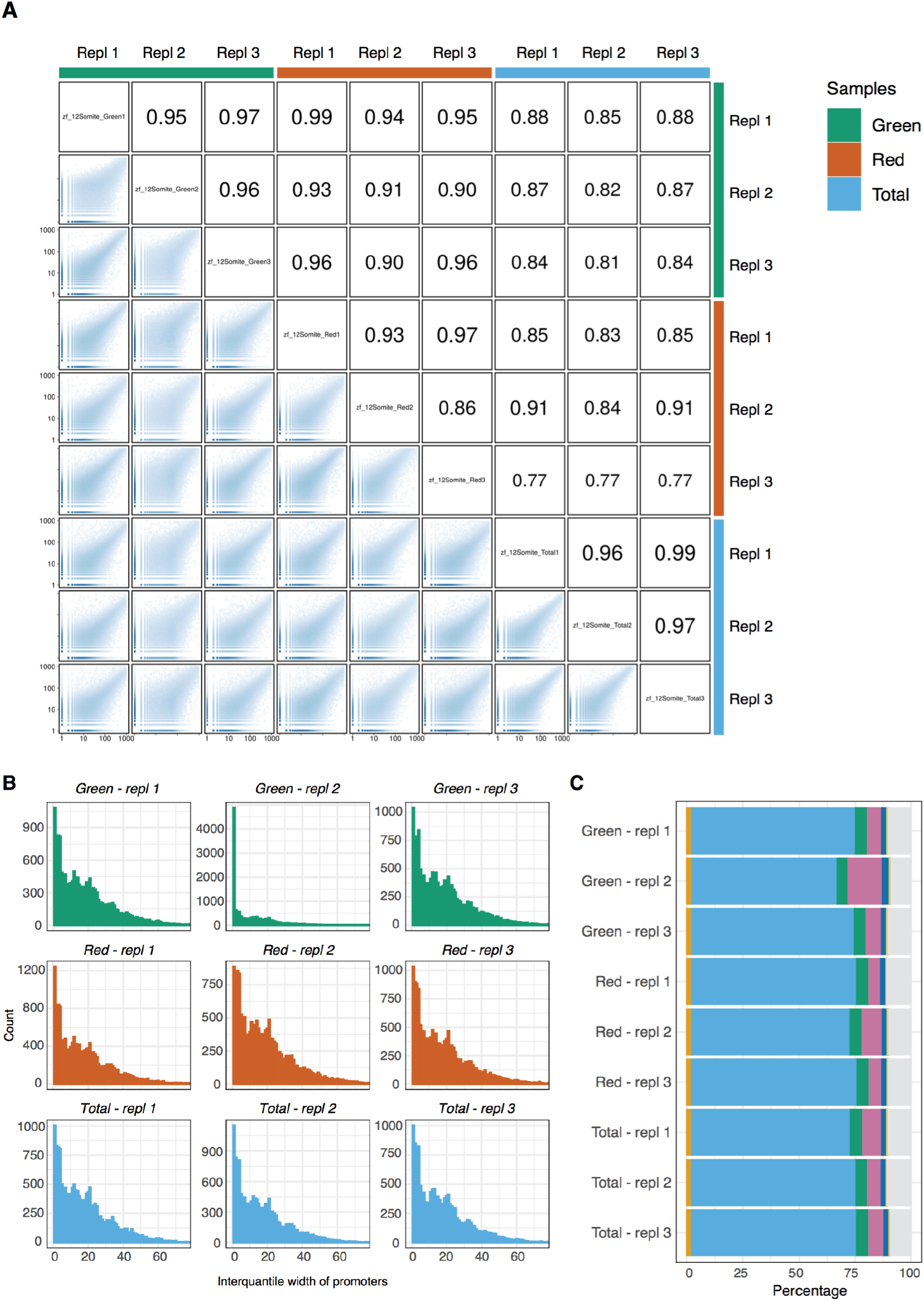
[A] CTSS correlation matrix for triplicate samples of G1 (Red), S/G2/M (Green) and unsegregated cells. [B] Graphs of tag cluster interquantile widths from the triplicate samples of G1 (Red), S/G2/M (Green) and unsegregated cells (Total). [C] Graph of mapping frequency to genomic features for triplicate samples of G1 (Red), S/G2/M (Green) and unsegregated cells. “Exon” (dark blue), “5’ UTR” (orange), “3’ UTR” (yellow) and “intron” (green) locations were extracted from the DanRer7 genomic build. “Promoter (<=1kb)” = window 0-1kb upstream of the gene start site (purple), “Promoter (1-3kb)” = 1-3kb region upstream of gene start (light blue), “Downstream <3kb” = window 0-3kb downstream of the gene end annotated in the DanRer7 genomic build (red), and “Distal intergenic” = all regions not covered in other classifications (grey).

**Supplementary figure 3.**
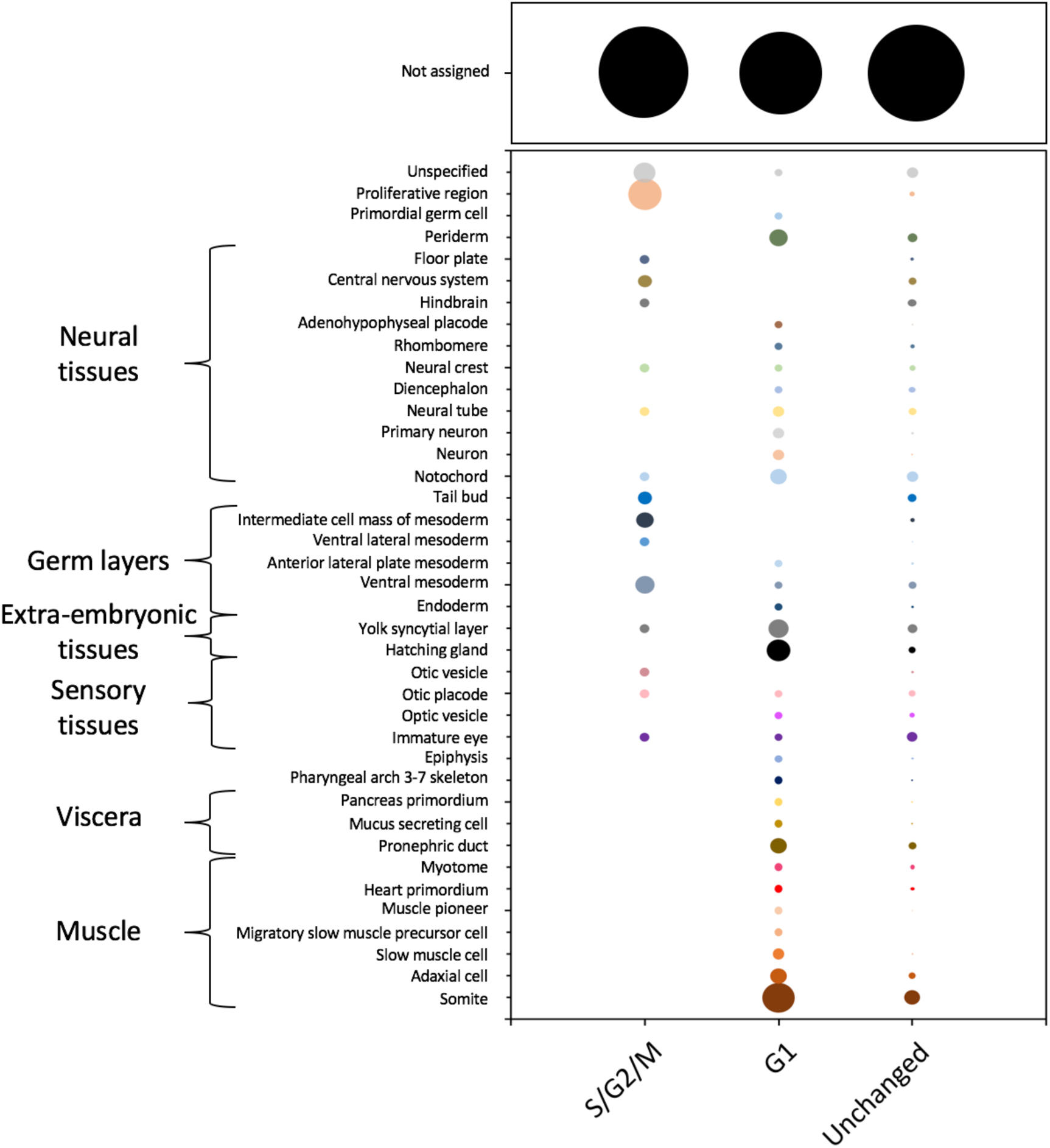
Bubble plot of tissue ontology of differentially expressed genes between S/G2/M and G1 cell populations (extracted from the ZFIN database of tissue specific gene expression for the 14-19 somite stage).

**Supplementary figure 4.**
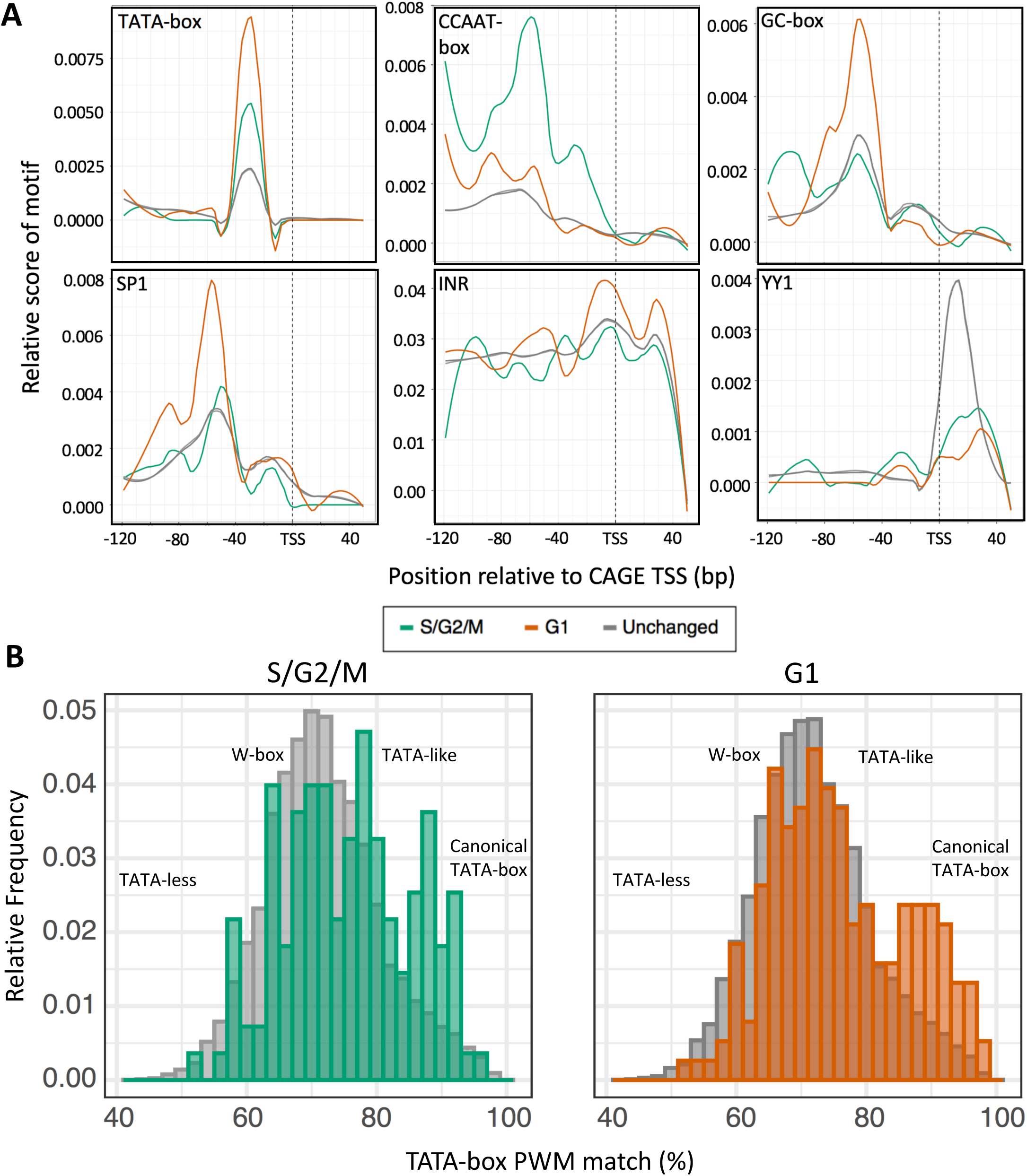
Additional core promoter architecture. [A} Metaplot of occurrence and positional constraint relative to TSS, of core promoter motifs (90% match) in different groups. [B] Distribution of position weight matrix (PWM) match (%) to TATA-box in the region –40 to –20 bp upstream of the dominant TSS in genes upregulated in S/G2/M (green) and G1 (orange) populations compared to genes with unchanged expression (grey).

**Supplementary figure 5:**
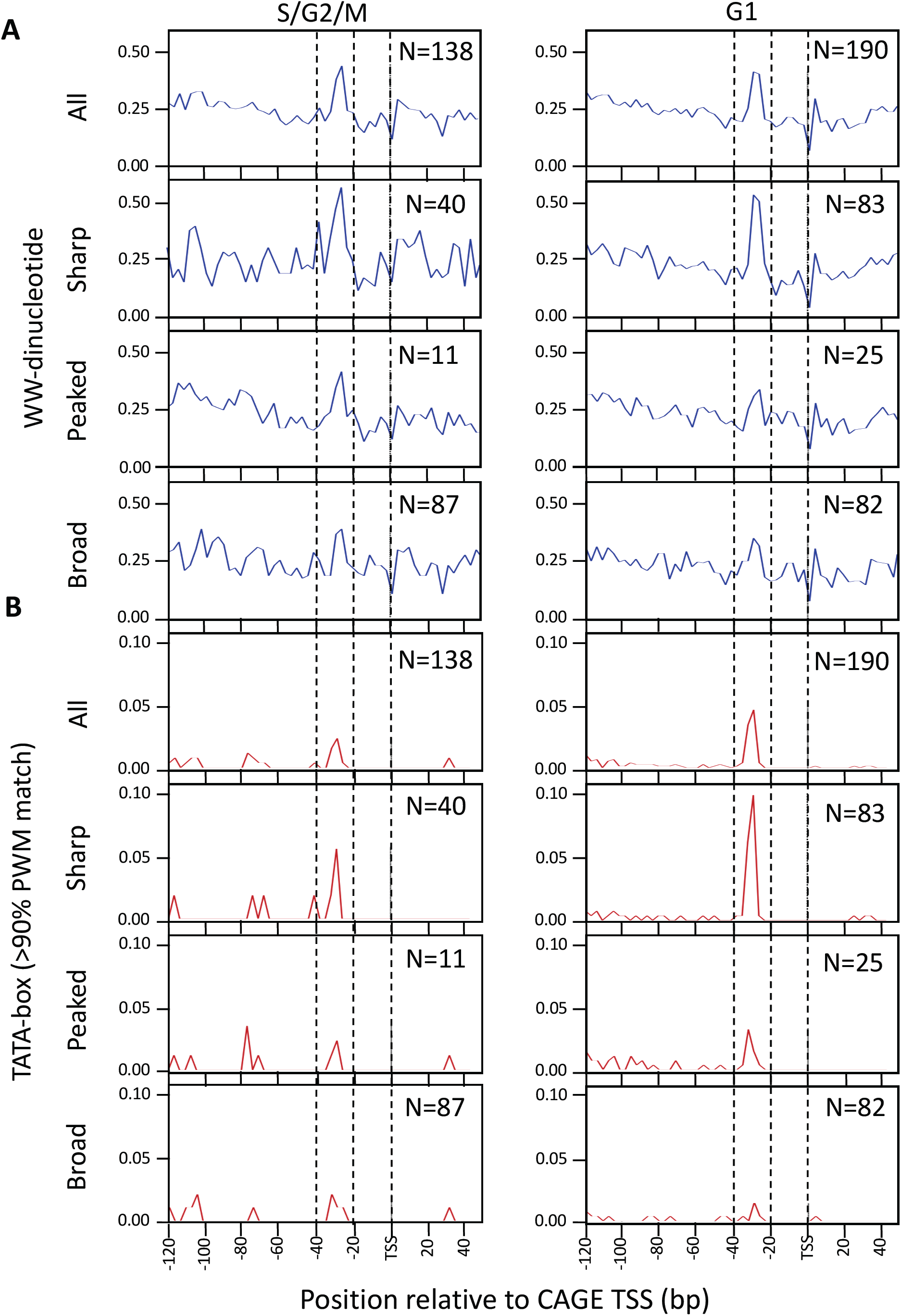
Analysis of differentially expressed gene promoter features. [A] WW Dinucleotide frequency analysis within the promoter region (120bp downstream and 50bp upstream of the dominant TSS) in genes with different promoter shapes, differentially expressed between G1 and S/G2/M. [B] TATA motif (>90% PWM match) frequency analysis within the promoter region (120bp downstream and 50bp upstream of the dominant TSS) in genes with different promoter shapes, differentially expressed between G1 and S/G2/M.

**Supplementary table 1:** Proportion of segregated cell displaying 2N-4N DNA content as determined by propidium iodide staining

|  | 2N | 2-4N | 4N |
| --- | --- | --- | --- |
| Total | 38.8 | 16.6 | 21.6 |
| Red (G1) | 80.9 | 2.3 | 6.7 |
| Green (S/G2/M) | 11 | 26.6 | 45.6 |

**Supplementary table 2:**
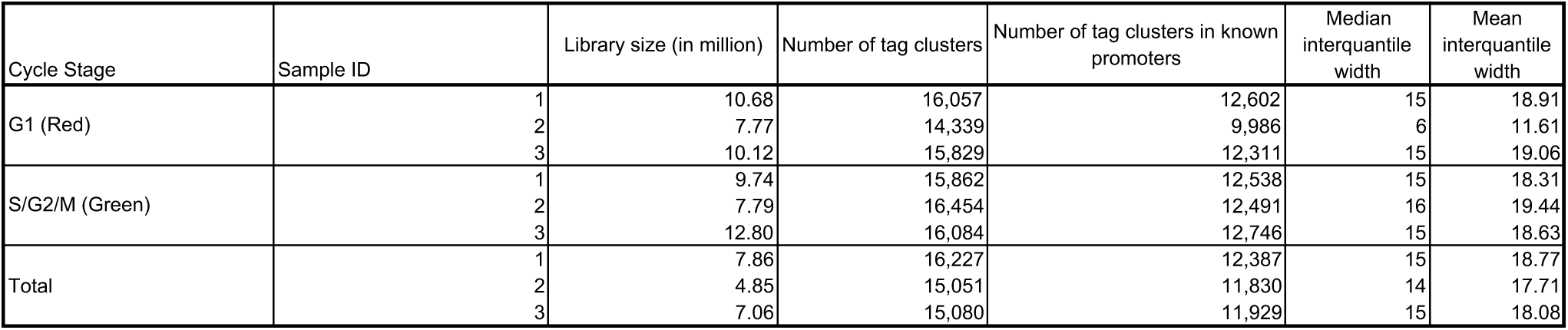
Key library and tag cluster statisitics for each sample replicate

**Supplementary table 3:** Key library and tag cluster statisitics for merged samples

| Cycle Stage | Library size (in million) | Number of tag clusters | Number of tag clusters in known promoters | IQ width (median) | IQ width (median) |
| --- | --- | --- | --- | --- | --- |
| 14_Somite_Red (G1) | 30.33 | 17,133 | 13,338 | 15 | 19.06 |
| 14_Somite_Green (S/G2/M) | 20.80 | 16,614 | 12,949 | 15 | 18.94 |
| 14_Somite_Total | 19.77 | 17,232 | 13,212 | 15 | 18.91 |
| 14_Somite_published_CAGE | 5.79 | 16,531 | 13,237 | 14 | 17.84 |

**Supplementary table 4:** TC overlap (%) between each merged sample and previously published CAGE

| sampleID | 14_Somite_Green(S/G2/M) | 14_Somite_Red(G1) | 14_Somite_Total | 14_Somite_published_CAGE |
| --- | --- | --- | --- | --- |
| 14_Somite_Green (S/G2/M) | 100.00 | 84.93 | 81.46 | 62.31 |
| 14_Somite_Red (G1) | 87.16 | 100.00 | 83.46 | 63.45 |
| 14_Somite_Total | 83.19 | 82.95 | 100.00 | 62.47 |
| 14_Somite_published_CAGE | 68.30 | 67.64 | 67.31 | 100.00 |

**Supplementary table 5:** GeneID overlap (%) between each merged sample and previously published CAGE

| sampleID | 14_Somite_Green(S/G2/M) | 14_Somite_Red(G1) | 14_Somite_Total | 14_Somite_published_CAGE |
| --- | --- | --- | --- | --- |
| 14_Somite_Green (S/G2/M) | 100 | 94.96249 | 94.15476 | 84.68378 |
| 14_Somite_Red (G1) | 97.13944 | 100 | 95.65303 | 85.95784 |
| 14_Somite_Total | 95.20582 | 94.55325 | 100 | 85.89368 |
| 14_Somite_published_CAGE | 92.08612 | 91.37679 | 92.37063 | 100 |

**Supplementary table 6:** Intersection between differentially expressed genes between samples and cell cycle periodic genes from Cyclebase

| de_group | Number of promoters | Number of human ortholog | Cell cycle base | Pvalue_permutation |
| --- | --- | --- | --- | --- |
| Green (S/G2/M) upregulated | 138 | 92 | 43 | < 1 e-4 |
| non-significant | 8406 | 6158 | 246 | n.s |
| red (G1) upregulated | 190 | 137 | 6 | n.s |

**Supplementary table 7:**
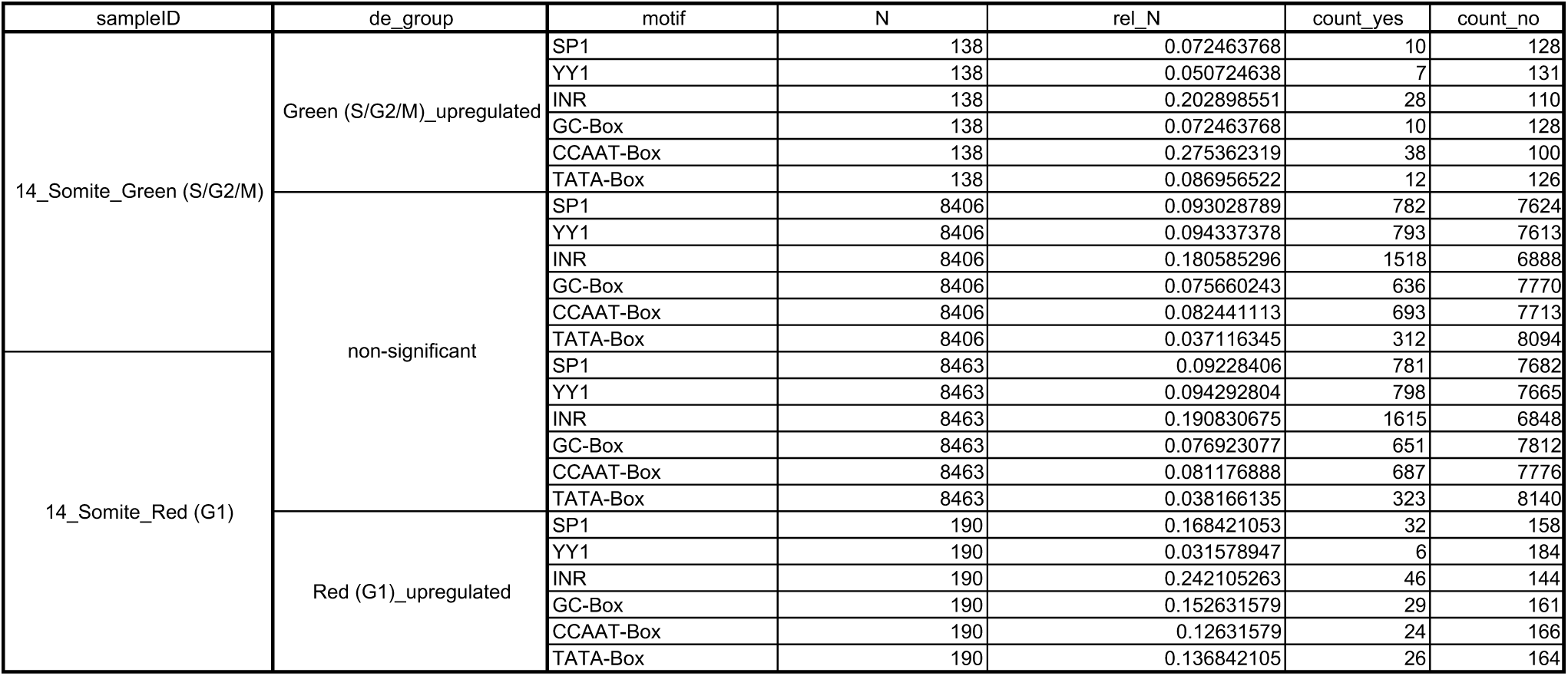
Promoter motif frequency analysis

**Supplementary table 8:**
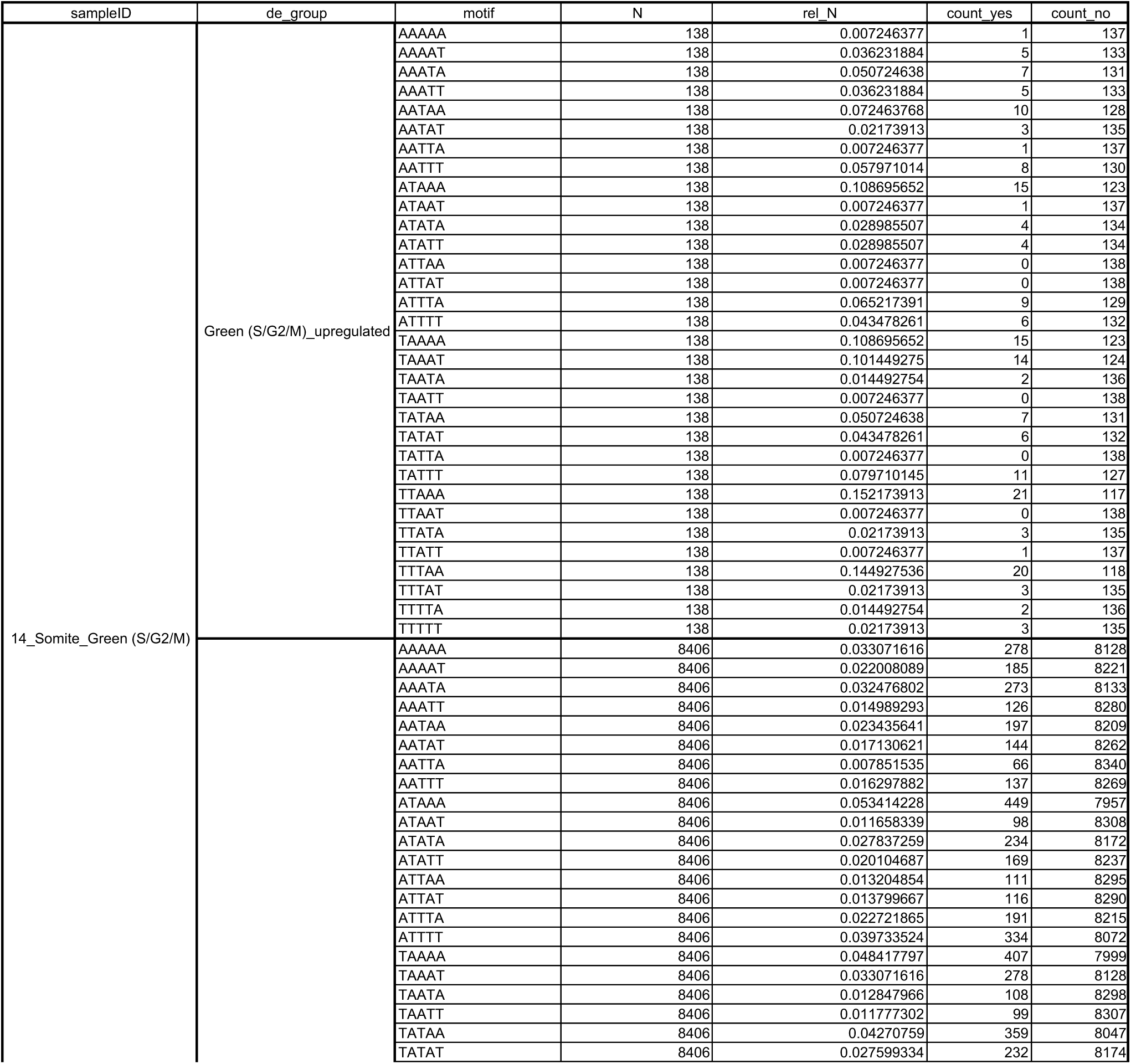
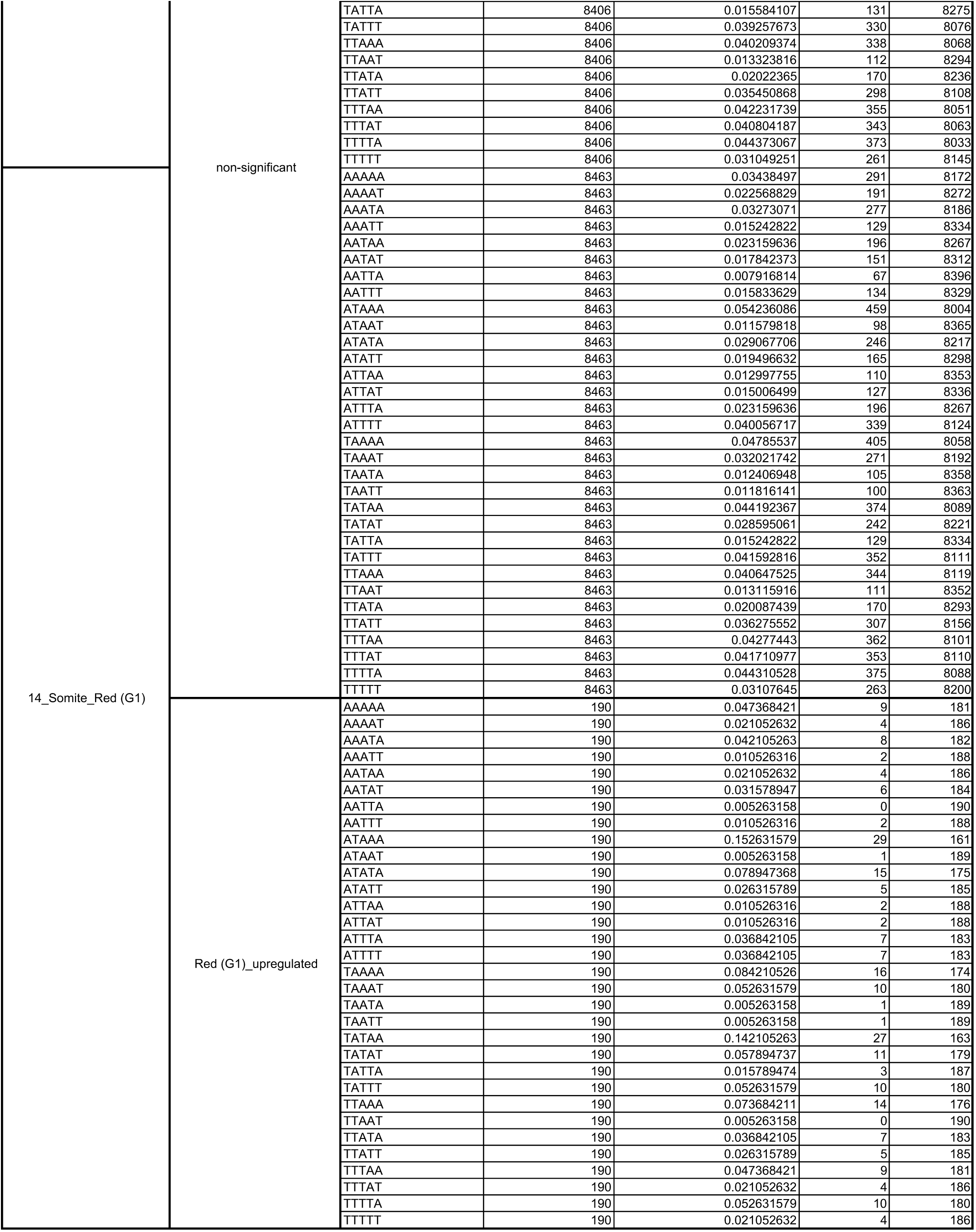
A/T pentamer frequency analysis

## References

1. D’Alessio JA, Ng R, Willenbring H, Tjian R. Core promoter recognition complex changes accompany liver development. Proc Natl Acad Sci U S A. 2011;108(10):3906–11.

2. Kadonaga JT. Perspectives on the RNA polymerase II core promoter. Wiley Interdiscip Rev Dev Biol. 2012;1(1):40–51.

3. Carninci P, Sandelin A, Lenhard B, Katayama S, Shimokawa K, Ponjavic J, et al. Genome-wide analysis of mammalian promoter architecture and evolution. Nat Genet. 2006;38(6):626–35.

4. Ponjavic J, Lenhard B, Kai C, Kawai J, Carninci P, Hayashizaki Y, et al. Transcriptional and structural impact of TATA-initiation site spacing in mammalian core promoters. Genome Biol. 2006;7(8):R78.

5. Sandelin A, Carninci P, Lenhard B, Ponjavic J, Hayashizaki Y, Hume DA. Mammalian RNA polymerase II core promoters: insights from genome-wide studies. Nat Rev Genet. 2007;8(6):424–36.

6. Lenhard B, Sandelin A, Carninci P. Metazoan promoters: emerging characteristics and insights into transcriptional regulation. Nat Rev Genet. 2012;13(4):233–45.

7. Akalin A, Fredman D, Arner E, Dong X, Bryne JC, Suzuki H, et al. Transcriptional features of genomic regulatory blocks. Genome Biol. 2009;10(4):R38.

8. Plessy C, Pascarella G, Bertin N, Akalin A, Carrieri C, Vassalli A, et al. Promoter architecture of mouse olfactory receptor genes. Genome Res. 2012;22(3):486–97.

9. Deaton AM, Bird A. CpG islands and the regulation of transcription. Genes Dev. 2011;25(10):1010–22.

10. Filipczyk AA, Laslett AL, Mummery C, Pera MF. Differentiation is coupled to changes in the cell cycle regulatory apparatus of human embryonic stem cells. Stem Cell Res. 2007;1(1):45–60.

11. Ohtsuka S, Dalton S. Molecular and biological properties of pluripotent embryonic stem cells. Gene Ther. 2008;15(2):74–81.

12. Calder A, Roth-Albin I, Bhatia S, Pilquil C, Lee JH, Bhatia M, et al. Lengthened G1 phase indicates differentiation status in human embryonic stem cells. Stem Cells Dev. 2013;22(2):279–95.

13. Roccio M, Schmitter D, Knobloch M, Okawa Y, Sage D, Lutolf MP. Predicting stem cell fate changes by differential cell cycle progression patterns. Development. 2013;140(2):459–70.

14. Kimmel CB, Ballard WW, Kimmel SR, Ullmann B, Schilling TF. Stages of embryonic development of the zebrafish. Dev Dyn. 1995;203(3):253–310.

15. Wragg J, Muller F. Transcriptional Regulation During Zygotic Genome Activation in Zebrafish and Other Anamniote Embryos. Adv Genet. 2016;95:161–94.

16. Haberle V, Li N, Hadzhiev Y, Plessy C, Previti C, Nepal C, et al. Two independent transcription initiation codes overlap on vertebrate core promoters. Nature. 2014;507(7492):381–5.

17. Nestorov P, Hotz HR, Liu Z, Peters AH. Dynamic expression of chromatin modifiers during developmental transitions in mouse preimplantation embryos. Sci Rep. 2015;5:14347.

18. Sugiyama M, Sakaue-Sawano A, Iimura T, Fukami K, Kitaguchi T, Kawakami K, et al. Illuminating cell-cycle progression in the developing zebrafish embryo. Proc Natl Acad Sci U S A. 2009;106(49):20812–7.

19. Deato MD, Tjian R. Switching of the core transcription machinery during myogenesis. Genes Dev. 2007;21(17):2137–49.

20. Muller F, Tora L. Chromatin and DNA sequences in defining promoters for transcription initiation. Biochim Biophys Acta. 2014;1839(3):118–28.

21. Haberle V, Stark A. Eukaryotic core promoters and the functional basis of transcription initiation. Nat Rev Mol Cell Biol. 2018;19(10):621–37.

22. Levine M, Cattoglio C, Tjian R. Looping back to leap forward: transcription enters a new era. Cell. 2014;157(1):13–25.

23. Westerfield M. The Zebrafish Book: A Guide for the Laboratory Use of Zebrafish (Danio Rerio): University of Oregon Press; 2000.

24. Poulain S, Kato S, Arnaud O, Morlighem JE, Suzuki M, Plessy C, et al. NanoCAGE: A Method for the Analysis of Coding and Noncoding 5’-Capped Transcriptomes. Methods Mol Biol. 2017;1543:57–109.

25. Kuhn RM, Karolchik D, Zweig AS, Wang T, Smith KE, Rosenbloom KR, et al. The UCSC Genome Browser Database: update 2009. Nucleic Acids Res. 2009;37(Database issue):D755–61.

26. Langmead B, Trapnell C, Pop M, Salzberg SL. Ultrafast and memory-efficient alignment of short DNA sequences to the human genome. Genome Biol. 2009;10(3):R25.

27. Haberle V, Forrest AR, Hayashizaki Y, Carninci P, Lenhard B. CAGEr: precise TSS data retrieval and high-resolution promoterome mining for integrative analyses. Nucleic Acids Res. 2015;43(8):e51.

28. Balwierz PJ, Carninci P, Daub CO, Kawai J, Hayashizaki Y, Van Belle W, et al. Methods for analyzing deep sequencing expression data: constructing the human and mouse promoterome with deepCAGE data. Genome Biol. 2009;10(7):R79.

29. Yu G, Wang LG, He QY. ChIPseeker: an R/Bioconductor package for ChIP peak annotation, comparison and visualization. Bioinformatics. 2015;31(14):2382–3.

30. Nepal C, Hadzhiev Y, Previti C, Haberle V, Li N, Takahashi H, et al. Dynamic regulation of the transcription initiation landscape at single nucleotide resolution during vertebrate embryogenesis. Genome Res. 2013;23(11):1938–50.

31. Adiconis X, Haber AL, Simmons SK, Levy Moonshine A, Ji Z, Busby MA, et al. Comprehensive comparative analysis of 5’-end RNA-sequencing methods. Nat Methods. 2018;15(7):505–11.

32. Takahashi H, Kato S, Murata M, Carninci P. CAGE (cap analysis of gene expression): a protocol for the detection of promoter and transcriptional networks. Methods Mol Biol. 2012;786:181–200.

33. Kawaji H, Lizio M, Itoh M, Kanamori-Katayama M, Kaiho A, Nishiyori-Sueki H, et al. Comparison of CAGE and RNA-seq transcriptome profiling using clonally amplified and single-molecule next-generation sequencing. Genome Res. 2014;24(4):708–17.

34. Carcamo J, Maldonado E, Cortes P, Ahn MH, Ha I, Kasai Y, et al. A TATA-like sequence located downstream of the transcription initiation site is required for expression of an RNA polymerase II transcribed gene. Genes Dev. 1990;4(9):1611–22.

35. Wang J, Xie Y, Bai X, Wang N, Yu H, Deng Z, et al. Targeting dual specificity protein kinase TTK attenuates tumorigenesis of glioblastoma. Oncotarget. 2018;9(3):3081–8.

36. Muller GA, Engeland K. The central role of CDE/CHR promoter elements in the regulation of cell cycle-dependent gene transcription. The FEBS journal. 2010;277(4):877–93.

37. Wasner M, Haugwitz U, Reinhard W, Tschop K, Spiesbach K, Lorenz J, et al. Three CCAAT-boxes and a single cell cycle genes homology region (CHR) are the major regulating sites for transcription from the human cyclin B2 promoter. Gene. 2003;312:225–37.

38. Wang W, Li Q, Bagchi IC, Bagchi MK. The CCAAT/enhancer binding protein beta is a critical regulator of steroid-induced mitotic expansion of uterine stromal cells during decidualization. Endocrinology. 2010;151(8):3929–40.

39. Ramskold D, Wang ET, Burge CB, Sandberg R. An abundance of ubiquitously expressed genes revealed by tissue transcriptome sequence data. PLoS Comput Biol. 2009;5(12):e1000598.

40. Fossat N, Pfister S, Tam PP. A transcriptome landscape of mouse embryogenesis. Dev Cell. 2007;13(6):761–2.

41. Farrell JA, Wang Y, Riesenfeld SJ, Shekhar K, Regev A, Schier AF. Single-cell reconstruction of developmental trajectories during zebrafish embryogenesis. Science. 2018;360(6392).

42. Kouno T, Moody J, Kwon AT, Shibayama Y, Kato S, Huang Y, et al. C1 CAGE detects transcription start sites and enhancer activity at single-cell resolution. Nat Commun. 2019;10(1):360.

43. FitzGerald PC, Sturgill D, Shyakhtenko A, Oliver B, Vinson C. Comparative genomics of Drosophila and human core promoters. Genome Biol. 2006;7(7):R53.

44. Ohler U. Identification of core promoter modules in Drosophila and their application in accurate transcription start site prediction. Nucleic Acids Res. 2006;34(20):5943–50.

45. Engstrom PG, Ho Sui SJ, Drivenes O, Becker TS, Lenhard B. Genomic regulatory blocks underlie extensive microsynteny conservation in insects. Genome Res. 2007;17(12):1898–908.

46. Rhee HS, Pugh BF. Genome-wide structure and organization of eukaryotic preinitiation complexes. Nature. 2012;483(7389):295–301.

47. Baptista T, Grunberg S, Minoungou N, Koster MJE, Timmers HTM, Hahn S, et al. SAGA Is a General Cofactor for RNA Polymerase II Transcription. Mol Cell. 2017;68(1):130-43.e5.

48. Watanabe K, Kokubo T. SAGA mediates transcription from the TATA-like element independently of Taf1p/TFIID but dependent on core promoter structures in Saccharomyces cerevisiae. PLoS One. 2017;12(11):e0188435.

49. Tapias A, Ciudad CJ, Roninson IB, Noe V. Regulation of Sp1 by cell cycle related proteins. Cell Cycle. 2008;7(18):2856–67.

50. Opitz OG, Rustgi AK. Interaction between Sp1 and cell cycle regulatory proteins is important in transactivation of a differentiation-related gene. Cancer Res. 2000;60(11):2825–30.

51. Grinstein E, Jundt F, Weinert I, Wernet P, Royer HD. Sp1 as G1 cell cycle phase specific transcription factor in epithelial cells. Oncogene. 2002;21(10):1485–92.

