## Supplementary movie legends for "Zebrafish embryonic tissue differentiation is marked by concurrent cell cycle dynamic and gene promoter regulatory changes"

**Supplementary movie 1:** Time-lapse fluorescent imaging of FUCCI embryo development, from high to 19 somite stage, imaged on the lightsheet microscope. Frames were acquired every 15 minutes.

**Supplementary movie 2:** Three dimensional rendering of a FUCCI embryo at the 14 somite stage. Imaged from the dorsal / posterior axis, on the lightsheet microscope.

**Supplementary movie 3:** Three dimensional rendering of a FUCCI embryo at the 14 somite stage. Imaged from the ventral/ anterior axis, on the lightsheet microscope.
